# Single-Cell Genomics Decontamination with CellSweep

**DOI:** 10.64898/2026.03.04.709349

**Authors:** Maya Caskey, Joseph Rich, Ryan Weber, Ali Mortazavi, Lior Pachter, Ingileif Hallgrimsdottir

## Abstract

Single-cell genomics technologies enable high-throughput cell profiling, but technical contamination remains an obstacle to accurate downstream analysis. Free-floating ambient molecules released from lysed cells and global bulk contamination introduced during library preparation can distort molecular profiles. These artifacts can obscure cellular identities and reduce the reliability of differential analysis or clustering results. We present an efficient and effective approach to removing ambient and bulk contamination that can be applied to data generated from a wide variety of technologies. We show that our tool, CellSweep, outperforms other methods to remove artifacts using numerous benchmarks.

## Introduction

Singe-cell genomics provides a fundamental framework for classifying cell types based on their molecular profile (Flynn et al., 2023). In order to achieve single-cell resolution, single-cell assays rely on the assignment of molecules to barcoded containers based on the physical or logical encapsulation of cells. Following library preparation, libraries are amplified and sequenced and individual transcripts are identified by their unique molecular identifiers (UMIs) and cell barcodes. Ideally, the contents of each barcoded container consist of molecules from a single cell. However, in practice, these assays are subject to several sources of error: ambient contamination from lysed or damaged cells introduces a background signal; containers may capture no or multiple cells (multiplets); and bulk contamination adds additional noise during amplification and sequencing (Luecken and Theis, 2019; Kavaliauskaite and Madsen, 2023; Griffiths et al., 2018; Farouni et al., 2020; Potapov and Ong, 2017). Notably, containers that capture no cell can still generate sequenced libraries composed entirely of background RNA. Although the physical mechanisms of cell partitioning differ between technologies, the resulting contamination can be described within a common statistical framework.

Sequencing technologies can be classified into three types according to the type of container: “shell”, “cell”, or “well” (Booeshaghi et al., 2023) (Fig. 1A). All three of these approaches suffer from ambient contamination, multiplets, and non-cellular barcodes, albeit to different degrees.

**Fig 1.**
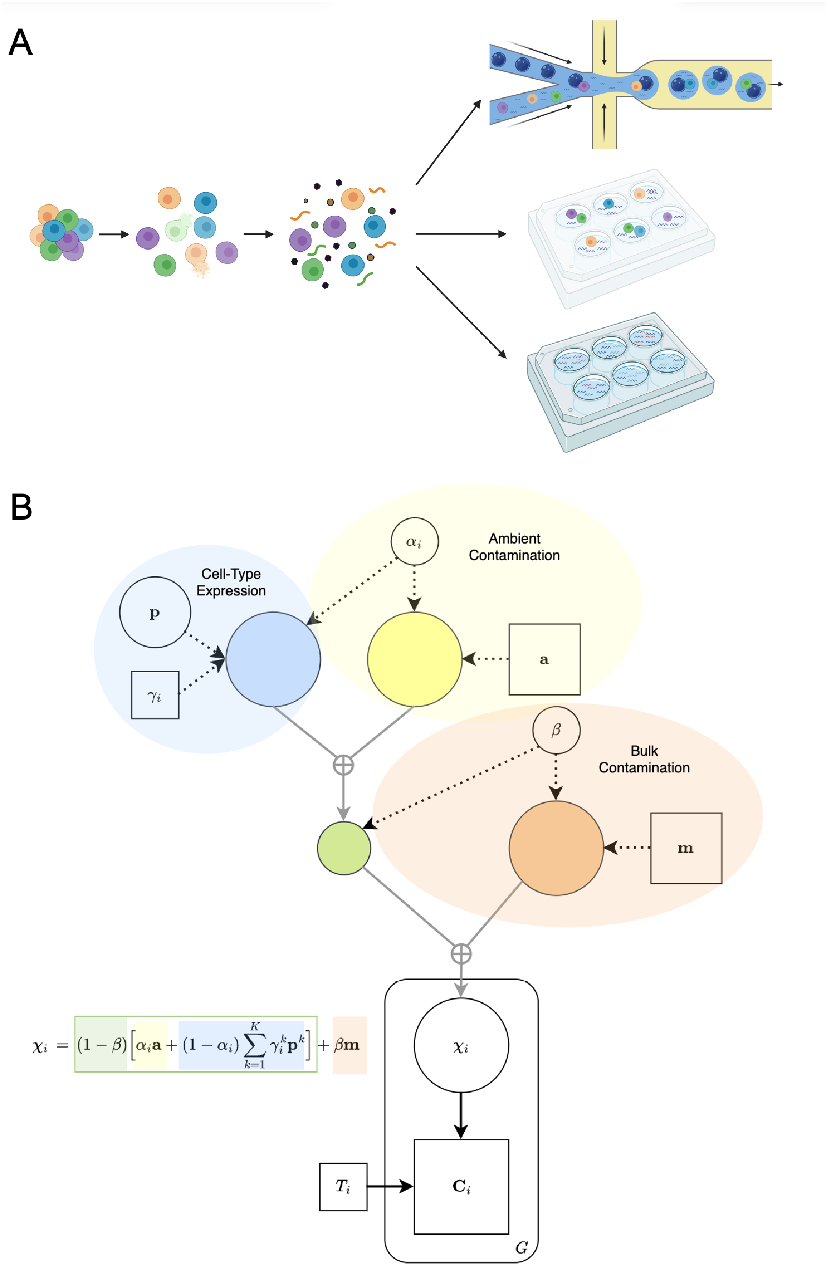
CellSweep overview. (A) Sources of ambient noise in “shell”/droplet technologies (top), “cell”/combinatorial barcode technologies (middle), and “well” technologies (bottom). (B) Diagram of the CellSweep model. Observed variables are shown as squares and latent variables as circles. The lower half shows the plate diagram for the multinomial generative model, and the upper half depicts the decomposition of ***χ***_*i*_ into ambient, cell-type, and bulk components.

We use the term “shells” to refer to the droplets used in droplet-based technologies such as 10x Genomics (Zheng et al., 2017), inDrops (Klein et al., 2015) and Drop-seq (Macosko et al., 2015). In these assays, a suspension of cells is partitioned into thousands of nanoliter droplets, each of which ideally contains both a single cell and a barcoded capture bead. After cell lysis, the bead binds and labels the RNA molecules. Ambient contamination arises from encapsulation of ambient RNA from the cell suspension along with a cell and a bead. To limit multiplets, these assays are intentionally underloaded, and a substantial portion of the recovered barcodes represent empty droplets that must be removed in downstream processing (Pan et al., 2022; Lun et al., 2019). This is similar for combinatorial barcoding technologies (Vitak et al., 2017).

Cells are the reaction chamber for combinatorial barcoding-based technologies such as SPLiT-seq (Rosenberg et al., 2018). In these assays, cells are labeled through successive rounds of combinatorial barcoding, in which cells are repeatedly partitioned into wells, assigned barcodes, and pooled. After multiple rounds, a RNA from a single cell can be identified by a unique permutation of barcodes. Although ambient RNA is not physically associated with an intact cell, it can still be captured and barcoded, producing ambient-only barcodes (equivalent to empty droplets) and contaminated cellular barcodes.

Technologies such as Smart-seq use wells to isolate single cells for amplification and sequencing (Ramsköld et al., 2012; Picelli et al., 2013). Ambient RNA molecules are introduced from the cell suspension prior to well partitioning. However, physical partitioning of cells into wells with fluorescent activated cell sorting significantly reduces instances of ambient contamination, non-cellular barcodes, and multiplets (Ding et al., 2020).

In addition to cell-level ambient contamination, bulk contamination can arise from molecular exchange during PCR amplification or sequencing (e.g. index hopping, barcode swapping, PCR chimeras) (Griffiths et al., 2018; Farouni et al., 2020; Potapov and Ong, 2017). This form of background noise affects all cells approximately uniformly and can be interpreted as a “global noise” profile superimposed on the true signal. Together, these contamination channels can substantially distort downstream analyses across single-cell technologies, leading to incorrect cell-type assignments, inflated cell–cell similarities, spurious marker expression, and compromised differential expression results.

Numerous computational methods have been developed to mitigate the effects background noise, particularly that introduced by ambient contamination, yet each comes with limitations. Methods relying on deep-generative models such as CellBender (Fleming et al., 2023) and scAR (Sheng et al., 2022) explicitly model contamination using variational inference and neural networks have a high computational cost: they often require GPUs and can take hours to run on medium-sized datasets. Other widely used tools—including DecontX (Yang et al., 2020) and SoupX (Young and Behjati, 2020)—are considerably faster, completing analyses in minutes on a CPU, but rely on simple generative models or heuristic corrections. The benchmarking that has been performed of these tools reveals divergent and variable performance, making it difficult to select tools in practice (Cargnelli et al., 2026).

To address these shortcomings, we have developed CellSweep, focusing on efficiency, accuracy and interpretability. The CellSweep model incorporates explicit mixture components for cell-type expression, ambient contamination, and global bulk contamination, and uses an efficient expectation—maximization (EM) algorithm for inference.

## Results

### Model

CellSweep models the observed counts corresponding to a given barcode as a mixture of three biologically interpretable sources: cell-type expression ***p***^*k*^, ambient contamination ***a***, and bulk contamination ***m*** (Fig. 1B), where ***p***^*k*^, ***a***, and ***m*** are vectors where each entry is the count assigned to that gene. We assume that all barcodes share a global bulk contamination fraction *β* and that each barcode *i* also has an individual ambient contamination fraction *α*_*i*_. For a cellular barcode *i*, with one-hot encoded cell-type assignment *γ*_*i*_, the expected expression distribution is:

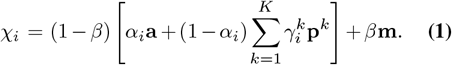

Conditioning on the total UMI count *T*_*i*_, we model the observed counts as:

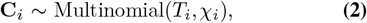

similar in spirit to the multinomial formulations in DecontX, SoupX, and scAR (Fig. 1B).

The CellSweep model assumes that cell-types are defined prior to inference with an arbitrary clustering or annotation method. Throughout this paper, we use CellTypist (Conde et al., 2022) to initialize the cell-type labels and determine the number of cell-types *K* unless otherwise specified.

Following approaches such as scAR and SoupX, CellSweep exploits the large number of non-cellular barcodes in typical datasets to obtain an empirical estimate of the ambient RNA profile. Since non-cellular barcodes contain only background RNA, their aggregated feature distribution provides a high-precision ambient estimate. After identifying non-cellular barcodes (e.g., via EmptyDrops or a hard threshold based on the knee-plot (Lun et al., 2019), CellSweep normalizes their counts to obtain the ambient profile ***a***, yielding a stable and unbiased estimator.

A central distinguishing feature of CellSweep is its use of the classical expectation—maximization (EM) algorithm for parameter inference (Dempster et al., 2018). In contrast to models such as CellBender, DecontX, and scAR, which rely on variational inference or deep generative architectures to approximate complex posteriors, CellSweep adopts a fully tractable generative likelihood that admits closed-form E- and M-steps. This yields substantial computational advantages: the E-step decomposes independently across cells and can therefore be parallelized with near-perfect scalability, while the M-step consists of simple normalized updates for the mixture components. As a result, the optimization procedure is both faster and more stable than variational approaches, without the need for stochastic gradient updates, neural-network architectures, or amortized inference. Although methods such as SoupX also avoid variational inference, their heuristics do not provide a unified probabilistic treatment of all sources of contamination. CellSweep thus achieves a balance of interpretability, computational efficiency, and modeling flexibility that distinguishes it from existing decontamination frameworks.

### An Alternative Model for Datasets without Non-Cellular Barcodes

The default CellSweep model assumes the presence of a sufficient number of non-cellular barcodes to obtain a reliable estimate of the ambient RNA profile. However, not all single-cell RNA-seq technologies produce non-cellular barcodes. In particular, well-based protocols such as Smart-seq2 may contain no non-cellular barcodes at all, yet still exhibit substantial ambient or bulk contamination. To broaden the applicability of CellSweep across scRNA-seq technologies, we introduce an alternative model that does not require non-cellular barcodes for inference.

In this alternative formulation, inspired by the approach used in DecontX, we assume that ambient background RNA arises from a mixture of cell-type expression profiles. Specifically, we model the ambient profile as a combination of the inferred cell-type profiles,

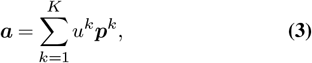

These mixture weights *u*^*k*^ are treated as latent parameters and are iteratively updated within the expectation—maximization (EM) framework.

This formulation is motivated by the observation that the ambient pool originates from lysed cells and therefore should approximate a mixture of cell-type expression profiles. However, it does not account for features that may be disproportionately represented in the ambient pool, such as mitochondrial transcripts released through organelle lysis (Gorin and Goodman, 2026).

### Background removal in mixed-species experiments

Background contamination in single-cell genomics data is readily identifiable in mixed-species experiments, when cells from two distinct species are processed and sequenced together. Ideally, all the reads associated with a barcode should originate from a single species. In practice, ambient contamination results in substantial off-target species counts.

We applied CellSweep and other methods to a publicly available human–mouse mixture dataset from 10x Genomics (Fig. 2). We removed doublets prior to analysis for all methods (see Supplementary Methods). This preprocessing step ensures that residual off-target counts primarily reflect ambient and bulk contamination rather than true biological mixtures. Most cells exhibit a small but meaningful amount of contamination, with a mean contamination of 1.25% for human cells and 2.93% for mouse cells. An ideal background-removal method would eliminate such cross-species counts while preserving same-species signal, although some removal of same-species counts is expected due to non–species-specific contamination.

**Fig 2.**
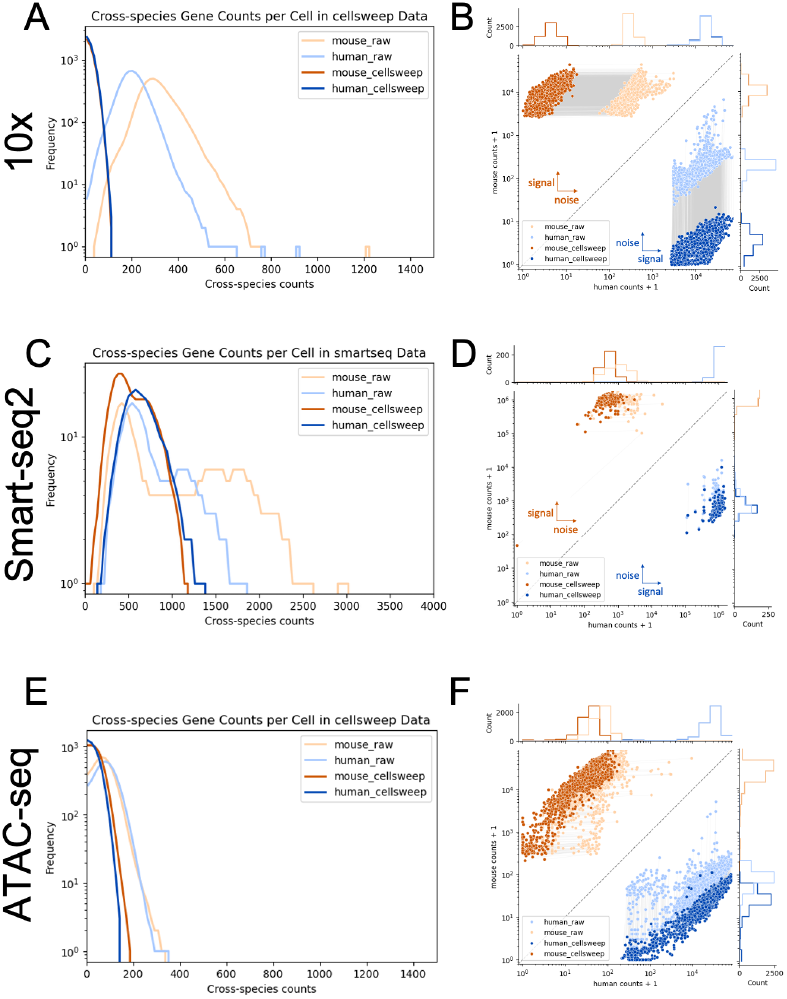
CellSweep effectively removes noise from human-mouse mixture data. (A) Histogram of total cross-species counts across all genes per cell after processing with CellSweep in a 10x dataset. (B) Joint scatterplot of mouse vs. human total counts per cell after processing with CellSweep in a 10x dataset. (C-D) Same as (A-B) in a Smart-seq2 dataset. (E-F) Same as (A-B) but in an ATAC-seq dataset. Light orange = mouse cells, raw; light blue = human cells, raw; dark orange = mouse cells, processed; dark blue = human cells, processed.

Fig. 2A-B summarizes the performance of CellSweep, and Fig. S1 summarizes the performance of SoupX, CellBender, DecontX, and scAR. For each approach, we show (i) the remaining cross-species counts per cells after denoising and (ii) the change in human and mouse UMI counts for each cell. Scatterplots describing the effect of each tool on the matrix entries, cells, and genes before vs. after processing are described in Fig. S2.

CellSweep removes cross-species contamination more consistently than all other methods tested (Fig. 2, Fig. S1). Across all cells, CellSweep reduces mouse gene contamination by 98.84% and human gene contamination by 98.59%, while retaining 97.85% of true-species counts in human cells and 98.46% in mouse cells. All other methods retain at least 97% of true species counts for both mouse and human, but fail to decontaminate all cells. CellBender and scAR substantially reduce cross-species contamination for most cells. CellBender removes 94.52% of mouse gene contamination and 97.29% of human gene contamination (Fig. S1C-D), while scAR removes 98.88% and 98.86%, respectively (Fig. S1G-H). However, both CellBender and scAR have a small handful of cells where over 90% of noise is retained. SoupX and DecontX remove relatively little cross-species contamination overall and display greater variability across cells. SoupX removes 87.97% of mouse gene contamination and only 66.65% of human contamination (Fig. S1A-B), while DecontX removes 81.57% and 68.25%, respectively (Fig. S1E-F). DecontX exhibits additional failures. After decontamination, DecontX retains cross-species noise directly proportional to the number of true-species counts, resulting in cells that appear to lie along a 45 degree line on the scatter plot. DecontX also removes nearly all counts from a small handful of cells and very little from others (Fig. S1F).

To further assess the performance of CellSweep, we computed the area under the curve (AUC) of off-target UMIs across cells for both mouse contamination in human cells and human contamination in mouse cells. In the raw data, these AUC values were 3,433,952.69 and 19,332,043.67, respectively. CellSweep reduced these values to 958.64 and 1279.02, which is substantially lower than those achieved by SoupX (159,153.52 and 392,445.66), Cell-Bender (136,480.40 and 456,481.74), DecontX (154694.48 and 11136.65), and scAR (83,604.30 and 110,136.32).

In addition to droplet-based technologies, the alternative CellSweep model can be used to remove noisy counts from well-based data. CellSweep reduces cross-species noise while retaining nearly all signal in a human-mouse mixture 685 cell scRNA-seq dataset generated with Smart-Seq 2 technology (Fig. 2C-D). On average, CellSweep retain 99.98% of signal in human cells and 99.80% of signal in mouse cells, while reducing human cell noise from 0.13% to 0.08%, and reducing mouse cell noise from 0.17% to 0.06%. Because of the nature of Smart-seq2 technology, many more counts are observed per cell compared to a typical cell from a 10x technology (on the order of 100,000 vs. 10,000 counts per cell, respectively), although both technologies have an average of approximately 1% ambient noise in these cells.

CellSweep is also useful for genomics assays other than scRNA-seq. We applied CellSweep to a 10X Genomics human-mouse mixture single-cell ATAC-seq dataset (Fig. 2E-F), and found that, just as with single-cell RNA-seq, CellSweep can reduce cross-species noise in ATAC-seq.

### Spatial transcriptomics

We ran CellSweep on a human/mouse Visium HD dataset in which cells from a human colorectal cancer cell line were grafted in a mouse. As with non-spatial droplet-based scRNAseq data, this dataset demonstrated a knee plot with a sharp inflection point, including hundreds of thousands of cells with fewer than 10 UMI counts (Fig 3A). After running CellSweep, much of the cross-species gene contamination is removed from human cells, with very little signal removed (Fig 3B). Most of the 249,802 cells have a predicted ambient noise fraction *α*_*i*_ near 0, although 40% of cells have a fraction over 0.1, and 9% of cells have a fraction over 0.5 (Fig. 3C). When visualizing predicted ambient noise fraction over spatial location, cells with a high *α*_*i*_ tend to lie along the edge of the tissue, indicating edge artifacts (Kummerfeld et al., 2025). In contrast, cells with a low *α*_*i*_ aggregated near the center of the tissue (Fig 3D).

**Fig 3.**
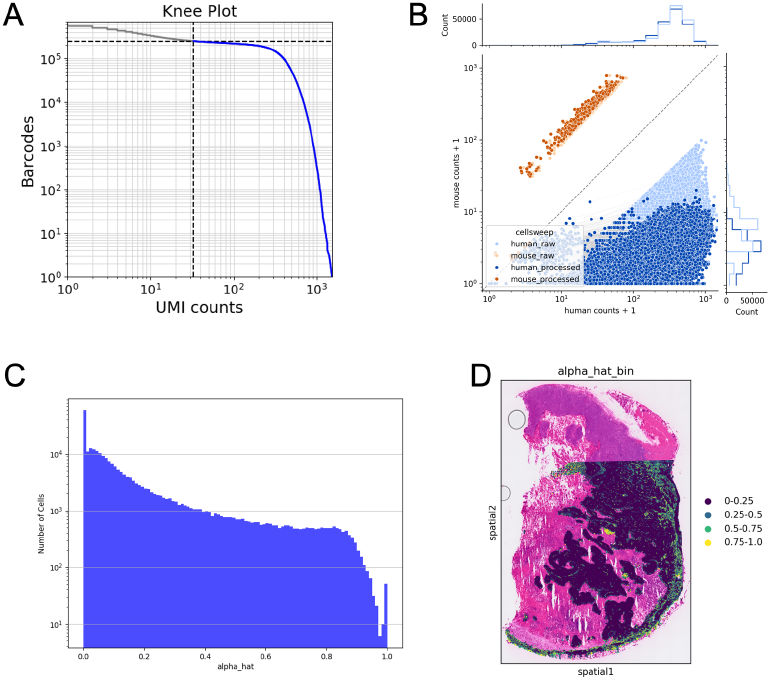
CellSweep predicts spatial localization of ambient noise in a human-mouse colorectal cancer xenograft Visium HD dataset. (A) Knee plot. (B) Joint scatterplot of mouse vs. human total counts per cell after processing with CellSweep. Light orange = mouse cells, raw; light blue = human cells, raw; dark orange = mouse cells, processed; dark blue = human cells, processed. (C) Histogram of alpha_hat (CellSweep’s predicted fraction of ambient noise per cell). (D) Spatial heatmap of binned alpha_hat values.

### Increased Cell-Type Marker Specificity in PBMC Data

Background contamination in scRNA-seq data reduces marker specificity and introduces spurious off-target expression that can propagate into downstream analyses (Janssen et al., 2023). To assess the impact of CellSweep and similar background-removal tools on biologically meaningful signal recovery, we evaluated performance on a publicly available 8,000-cell PBMC dataset, following a benchmarking strategy similar to that of Fleming et al. (2023) (Figs. 4, -S6).

**Fig 4.**
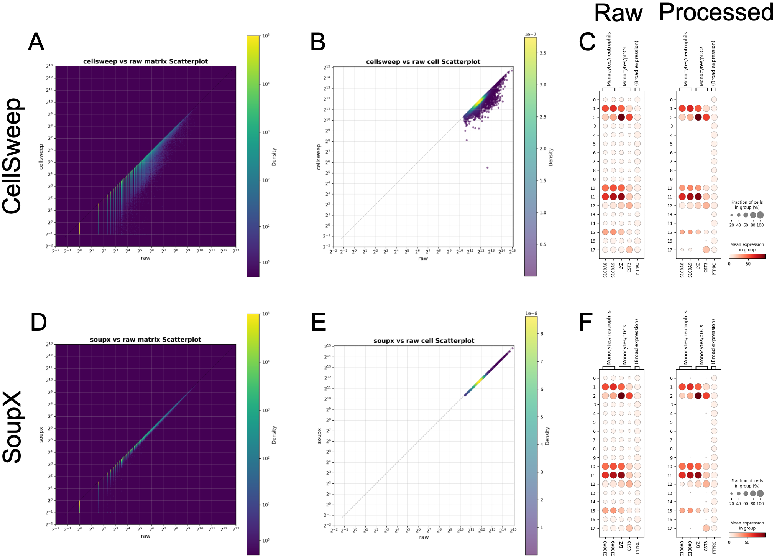
CellSweep effectively removes noise from a human PBMC 8k dataset. (A) Scatterplot of matrix values after vs. before processing with CellSweep. (B) Scatterplot of total cell counts after vs. before processing with CellSweep. (C) Dotplots of markers from monocytes/neutrophils, monocytes/pDCs, and broad expression in raw (left) and processed (right) data with CellSweep. (D-F) Same as (A-C) but with SoupX.

Prior to background correction, several well-known immune marker genes exhibit broad, non-specific expression across clusters. In particular, S100A8, S100A9, LYZ, CST3, and PTPRC appear at appreciable levels in nearly all clusters (Fig. 4C, left). However, S100A8 and S100A9 are canonical neutrophil markers, while LYZ and CST3 are primarily associated with monocytes and plasmacytoid dendritic cells. Their ubiquitous expression in the uncorrected data therefore reflects technical contamination rather than true biological signal. All tools succeed in removing counts of monocyte/dendritic cell/neutrophil marker genes from clusters that likely represent other cell types (Fig. 4C,F, S3C, S3F, S3I). In contrast, PTPRC, a pan-leukocyte marker that is expected to be broadly expressed across immune cell types, is retained uniformly across clusters after denoising.

In our benchmark, each tool removes counts to different degrees. The fewest mean counts per cell removed was by CellBender at 121.29 counts, followed by SoupX at 315.45, CellSweep at 667.88, DecontX at 770.32, and scAR at 2,647.38. Notably, because of its conservative removal of counts, CellBender appears to have been the least successful at eliminating marker contamination. SoupX was the only tool that did not noticeably alter any cells in terms of total counts compared to the raw matrix (Fig.4E). Overall, scAR was the most agressive denoiser and removed more counts per cell than any other tool (Fig. S3H).

Comparison of the changes in counts to matrix, total cell, and total gene counts between CellSweep and other tools reveals that all tools share similarities in their output (Fig. S5). In descending order of similarity with other tools, CellSweep most resembles DecontX, followed by SoupX, CellBender, and finally scAR. scAR consistently removes more counts in nearly all cells compared to CellSweep; all other tools do not possess any considerable trends.

### Noise reduction in a complex multiplexed experiment

We applied CellSweep to the 8 cubed founder and Trem2 datasets. These datasets serve as valuable controls because, unlike the previously-analyzed datasets, they (1) were generated with the Parse Biosciences Evercode WT v2 assay (i.e., non-droplet), (2) are derived exclusively from mouse (i.e., non-human), (3) involve eight different tissues, (4) are larger and more complex in scale (600,000-900,000 cells per plate), and (5) demonstrate well-characterized noise (Rebboah et al., 2026).

The 8 cubed founder experiment is designed so that each plate contains two different tissues, allowing for more accurate downstream quantification of batch effects and plate-specific contamination (Fig. 5A). CellSweep almost entirely eliminates cross-tissue marker contamination while retaining tissue-specific markers. We present two representative examples of this result with the cortex/hippocampus marker Snap25 on plate igvf_003 (Fig. 5B-C) and the adrenal marker Star on plate igvf_009 (Fig. 5D-E).

**Fig 5.**
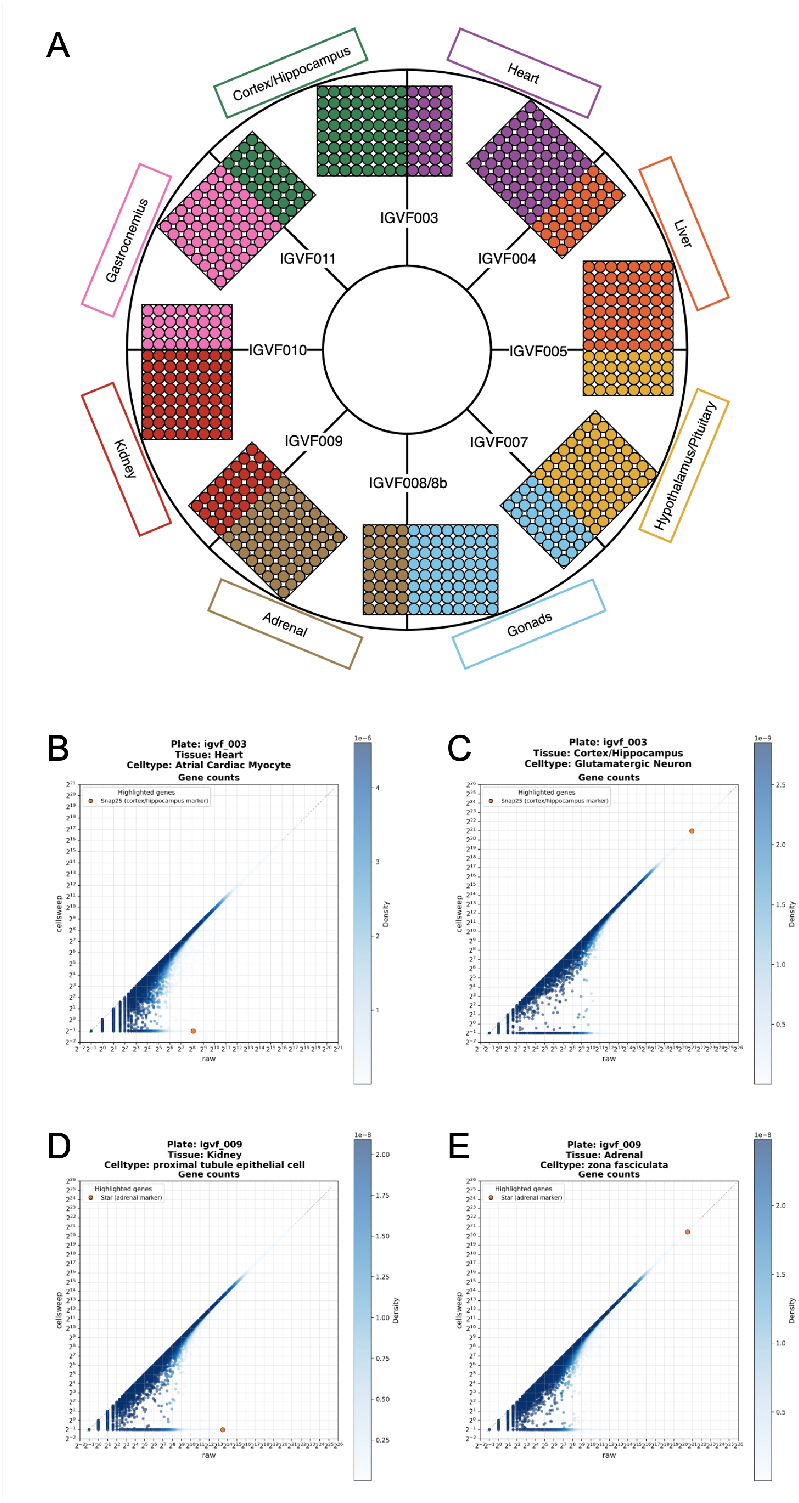
Cellsweep reduces cross-tissue contamination from marker genes in 8 cubed founder data. (A) Schematic of 8 cubed founder scRNA-seq setup. Color = tissue. (B) Scatterplot of total gene counts of Snap25 (a cortex/hippocampus marker) in processed vs. raw cells with CellSweep in atrial cardiac myocyte heart cells from plate igvf_003 (cortex/hippocampus and heart). (C) Scatterplot of total gene counts of Snap25 (a cortex/hippocampus marker) in processed vs. raw cells with CellSweep in glutamatergic neuron cortex cells from plate igvf_003 (cortex/hippocampus and heart). (D) Scatterplot of total gene counts of Star (an adrenal marker) in processed vs. raw cells with CellSweep in proximal tubule epithelial kidney cells from plate igvf_009 (adrenal and kidney). (E) Scatterplot of total gene counts of Star (an adrenal marker) in processed vs. raw cells with CellSweep in zona fasciculata adrenal cells from plate igvf_009 (adrenal and kidney).

The Trem2 dataset, which contains 161,200 cells from all eight tissues sequenced on a single plate, serves as an additional control. In this data set, CellSweep reduces the count of the liver marker albumin in all non-liver tissues while leaving counts in the liver relatively intact. CellSweep demonstrates similar success on the Trem2 dataset with the muscle markers Myh4 and titin (Fig. S7).

### Idempotency

As a control, we investigated the performance of programs when run repeatedly on the same dataset. Ideally, once a dataset is denoised, further denoising should not identify new noise, i.e. tools should be idempotent. Idempotency indicates model stability and ensures that cleaned data are not progressively eroded with repeated application.

We evaluated idempotency for CellSweep and other methods by reapplying each tool three additional times to the 8,000-cell PBMC dataset after an initial denoising step. As shown in Fig. 6 and Fig. S8, CellSweep, SoupX, and DecontX demonstrate near idempotency, with only minor changes observed after the second iteration. In contrast, Cell-Bender removes additional counts upon reapplication, and scAR continues to alter thousands of cells even after two iterations, indicating less stable behavior.

**Fig 6.**
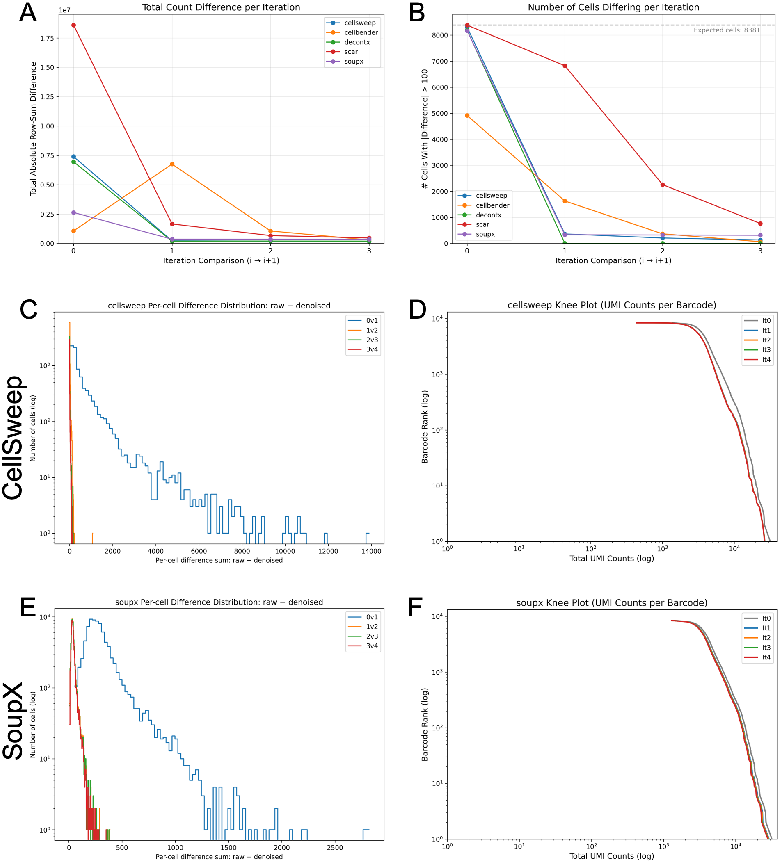
CellSweep demonstrates idempotency. (A-B) Total count difference (A) and number of cells differing by more than 100 counts (B) across iterations. Points have been connected for ease of visualization only (i.e., no interpolation). Blue = CellSweep; orange = CellBender; green = DecontX; red = scAR; purple = SoupX. (C) Histogram of per-cell count differences after processing with CellSweep. Blue = between iteration 0 (raw) and 1; orange = between iteration 1 and 2; green = between iteration 2 and 3; red = between iteration 3 and 4. (D) Knee plots after processing with CellSweep. Gray = iteration 0 (raw). Blue = iteration 1; orange = iteration 2; green = iteration 3; red = iteration 4. (E-F) Same as (C-D) but with SoupX. Analysis performed on the PBMC 8k dataset.

In the event that noncellular barcodes are unavailable, the alternative CellSweep model demonstrates a loss in idempotency (Fig. S8G-H).

### Runtime

CellSweep exhibits faster runtimes than existing background-removal methods, able to run on full datasets in under a minute. Fig. 7 summarizes the runtime of each method on the 8,000-cell PBMC dataset. When run on a single CPU thread, CellSweep completes in approximately 5 minutes. Using 16 CPU threads reduces runtime to 10 times faster at 25 seconds, making CellSweep over twice as fast as DecontX and SoupX under comparable conditions. Switching to the alternative CellSweep model in the absence of non-cellular barcodes does not appreciably change runtime. In contrast, neural-network-based methods such as CellBender and scAR require substantially longer runtimes, on the order of hours on CPU and between 10-30 minutes on GPU.

**Fig 7.**
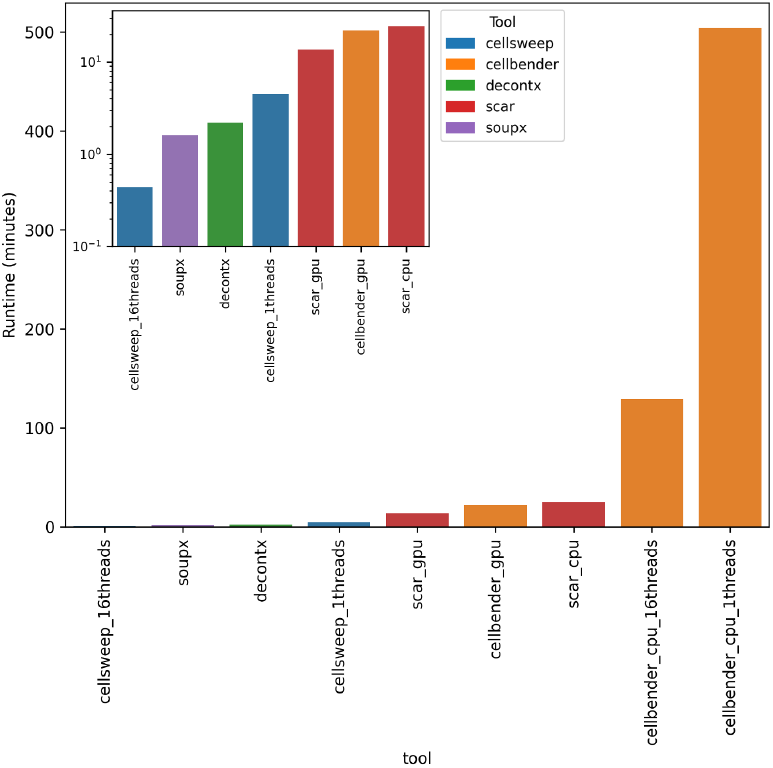
Runtime analysis. Analysis performed on the PBMC 8k dataset. Inset = log scale with tools under 30 minutes. Blue = CellSweep; orange = CellBender; green = DecontX; red = scAR; purple = SoupX.

### Simulation

In order to test these tools on a dataset with a clearly-defined ground truth, we developed a simulation of an scRNA-seq experiment, and generated a count matrix for a 10,000 gene, 10,000 cell dataset with 90,000 non-cellular barcodes and 12 artificially constructed cell-types (Fig. 8A). Simulated data were generated assuming negative binomial–distributed cell-type expression and Poisson-distributed ambient noise, consistent with both empirical observations and established probabilistic models of scRNA-seq count data (Hafemeister and Satija, 2019; Gorin and Goodman, 2026; Fleming et al., 2023). Bulk noise was additionally modeled as a random redistribution of counts across cells (see Supplementary Methods for additional details). Compared to real datasets, the simulated data exhibit little contamination, with a global bulk contamination of 5% and a median ambient contamination of 3%. Using the expression of the 20 most contaminating cell-type markers across cell-types, the dotplot of the raw data shows clear marker expression and little marker contamination (Fig. 8B).CellSweep, SoupX, CellBender, and DecontX all effectively remove noise while removing minimal signal (Fig. 8C-F, Fig. S12A-D, Fig. S13). scAR, on the other hand, fails to decontaminate the simulated data. Instead, it adds noisy counts and removes signal, as can be seen by the disappearance of expression of genes 6031, 4646, and 927 in cell type 4; 6544 in cell type 8; and 5794 in cell type 10 (Fig. S12E-F). Moreover, scAR has the lowest positive predictive value (PPV) at 0.67, indicating that scAR aggressively removes true signal across the dataset. All other tools have a PPV of 0.98 or greater. CellSweep was effective at removing off-target counts of marker genes while retaining on-target counts, reducing the median number of off-target counts per cell 23 to 3.85, while only reducing median signal from 733 to 730.53 (Fig. 8C-D). SoupX, CellBender, and DecontX performed similarly well, reducing noisy counts per cell from 23 to 4.02, 3.00, and 3.47, respectively (Fig. 8E-F, Fig. S12A-D).

**Fig 8.**
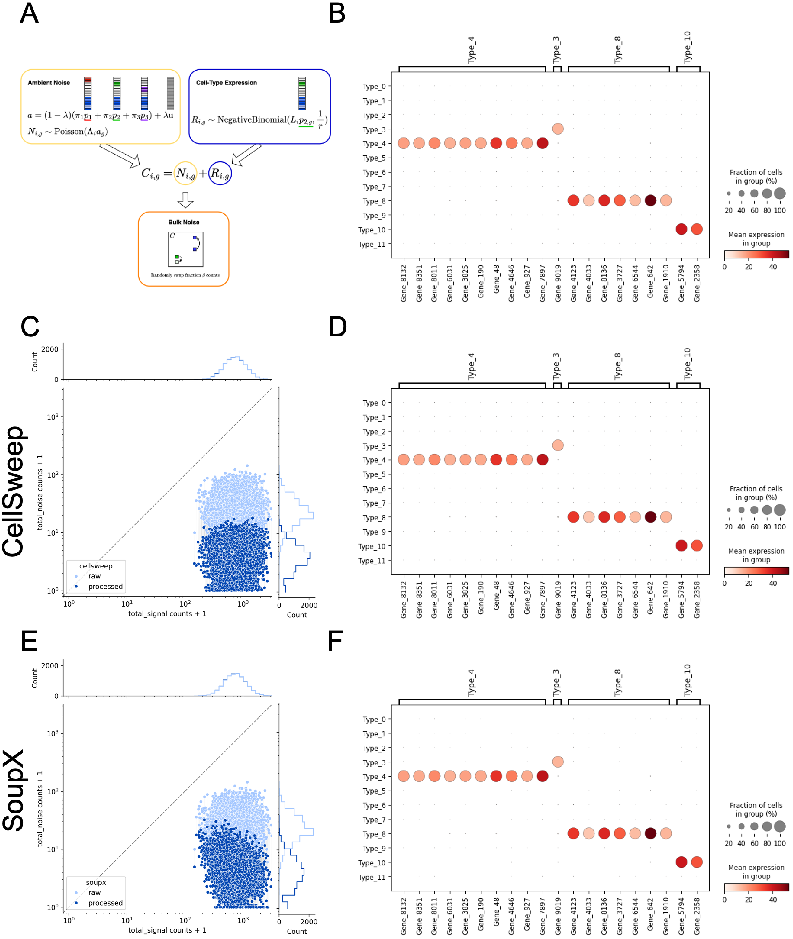
CellSweep removes noise in simulation. (A) Diagram of model for simulated dataset, assuming three cell-types (1 (red), 2 (green), and 3 (purple)) and cell-type assignment of 2 to cell *i*. Striped boxes represent the artificially constructed cell-type profiles *p*_*k*_ with red, green, and purple indicating the respective cell-type markers and blue boxes indicating house-keeping genes. Varying color intensities illustrate varying levels of gene expression. The gray box represents a uniform distribution of counts across all genes. The simulation assumes that a count matrix is the sum of Poisson distributed noise and negative-binomial distributed cell-type expression. Bulk noise is introduced by random movement of counts from one cell to another. B) Dotplot of the twenty most highly contaminating marker genes across all cell-types in the simulated data. (C) Joint scatterplot of noise and signal total marker counts per cell after processing with CellSweep. Light blue = raw; dark blue = CellSweep. (D) Same as (B) but after denoising with CellSweep. (E-F) Same as (C-D) but with SoupX.

## Discussion

Ambient and bulk contamination systematically distort scRNA-seq measurements and can compromise biological interpretation. Existing correction strategies often trade modeling rigor for computational efficiency, leaving a gap between deep generative approaches and lightweight heuristic methods. CellSweep bridges this gap by providing a fast, interpretable generative framework for routine decontamination. CellSweep adopts an explicit probabilistic model whose parameters correspond directly to biologically meaningful quantities. Inference proceeds via the EM algorithm over mixture components representing cell-type expression, ambient RNA, and global bulk contamination. Unlike deep generative models such as CellBender and scAR, which rely on neural networks and variational inference, CellSweep performs direct likelihood-based optimization. This formulation provides both interpretability and substantial computational advantages: CellSweep runs in minutes on a CPU for typical datasets and requires no specialized hardware.

Beyond denoising, CellSweep provides a per-cell estimate of ambient contamination, 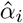 which serves as a quantitative indicator of cell quality. In practice, this metric identifies lowquality or noisy cells and clusters not reliably detected by conventional filtering heuristics such as UMI thresholds or mitochondrial content. Owing to its speed, minimal hyperparameter tuning required, and low sensitivity to initialization choices, CellSweep can be applied robustly across diverse datasets with minimal user intervention. These properties support treating ambient and bulk correction as a routine preprocessing step rather than a specialized adjustment.

Across a range of technologies, species, cell counts, and experimental conditions—including droplet-based, well-based, spatial, and single-nucleus datasets—CellSweep consistently improved marker specificity and reduced cross-sample contamination. Sensitivity analyses demonstrated robustness to clustering resolution, noncellular barcode identification strategy, and subsampling of noncellular barcodes (Fig. S9-11). Even under aggressive subsampling, results remained stable until 10,000 noncellular barcodes, supporting the practical reliability of the method across preprocessing pipelines (Fig. S11). If a dataset has fewer than 10,000 noncellular barcodes, we recommend using our alternative CellSweep model instead.

Future directions include both methodological extensions and downstream applications. On the modeling side, incorporating doublet detection directly into the generative framework would enable joint inference of contamination and multiplet structure, particularly in dense droplet-based experiments. Further improvements to the alternative ambient-learning regime to recover idempotency and increase statistical power will enhance robustness on datasets lacking non-cellular barcodes.

An especially important direction is the systematic study of how ambient and bulk decontamination influence down-stream machine learning models. Large-scale single-cell atlases are increasingly used to train foundation models, representation learning frameworks, and predictive classifiers. Because these models are sensitive to structured noise, ambient contamination may be implicitly encoded in learned embeddings, decision boundaries, and inferred regulatory programs. Applying CellSweep prior to model training provides an opportunity to quantify how removing structured technical signal alters learned representations, improves predictive performance, and enhances biological interpretability. Ongoing work in our group focuses on retraining and benchmarking machine learning models on CellSweep-processed datasets to assess the extent to which decontamination changes model behavior at scale.

**Table 1.** Comparison of scRNA-seq denoising tools.

| Metric | CellBender | DecontX | scAR | SoupX | CellSweep |
| --- | --- | --- | --- | --- | --- |
| Human-Mouse denoising | ✓ | X | ✓ | X | ✓ |
| PBMC Marker Cleanup (dotplots) | ✓ | ✓ | ✓ | ✓ | ✓ |
| Idempotent | X | ✓ | ✓ | ✓ | ✓ |
| Runtime on CPU (min) | 180 | 3 | 200 | 2 | 1 |
| Simulation PPV | 0.985 | 0.981 | 0.686 | 0.980 | 0.981 |
| Doesn't predict bigger counts | ✓ | ✓ | X | ✓ | ✓ |

Together, these directions position CellSweep not only as a preprocessing tool, but as a foundation for more reliable statistical and machine learning analyses of single-cell data.

In summary, CellSweep provides a fast, interpretable, and broadly applicable solution for separating biological signal from structured technical noise in single-cell data. As datasets continue to grow in size and complexity, methods that combine statistical transparency with computational efficiency will be essential, and CellSweep offers a practical foundation for this goal.

## Data Availability

- **hgmm_12k**: 10x hgmm_12k
- **kidney_nuclei_10k**: 10x Kidney Nuclei 10k
- **pbmc_mouse_5k**: 10x Mouse PBMC 5k
- **pbmc8k**: 10x PBMC 8k
- **pbmc33k**: 10x PBMC 33k
- **melanoma**: 10x Melanoma 10k
- **8-cubed founder strains** (Rebboah et al., 2026):

IGVF 8-Cubed SPLiT-seq

- **ATAC-seq**: 10x ATAC Mixture
- **Smart-seq2**: GSE132044 (GEO)

## Code Availability

The CellSweep Python package is available at https://github.com/pachterlab/cellsweep, along with notebooks to reproduce all analysis in this study.

## Author Contributions

Conceptualization, I.H, motivated by observations of R.W. and A.M. Methodology, M.C., J.R., L.P., and I.H. Investigation, L.P. and I.H. Visualization, M.C., J.R., L.P., and I.H. Funding acquisition, A.M., L.P., and I.H. Project administration and supervision, L.P. and I.H.. Writing—original draft, M.C. and J.R. Writing—review and editing, M.C., J.R., R.W., A.M., L.P., and I.H.

## Acknowledgments

We thank the Pachter Lab for helpful feedback at the start of the project. The project was funded by the Caltech Bioinformatics Resource Center (CBRC) and the Impact of Genomic Variation on Function (IGVF) Consortium under award number UM1HG012077. Thanks to the CBRC and the IGVF consortium for financial and resource support for this project.

## Methods

Given a barcode by feature count matrix, the CellSweep mixture model includes parameters for cell-type expression, ambient contamination, and bulk contamination, in a fully generative framework. Each barcode *i* produces a count vector

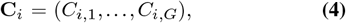

with total counts *T*_*i*_ = Σ_*g*_ **C**_*i,g*_. Conditional on its total count, we assume that

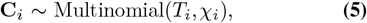

where 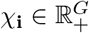represents the expected normalized feature expression profile of barcode *i*.

### Mixture Composition

We model *χ*_*i*_ as a convex combination of three biologically interpretable sources:

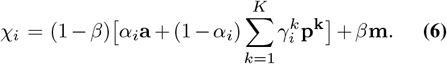

Equivalently,

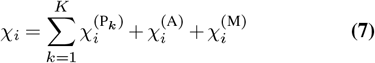

where,

- 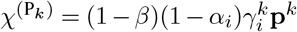is the contribution from cell-type *k*;
- *χ*^(A)^ = (1−*β*)*α*_*i*_**a** is the contribution from ambient contamination;
- *χ*^(M)^ = *β***m** is the contribution from bulk contamination.

Here,

- **a***∈*ℝ^*G*^ is the ambient contamination profile, representing molecular contamination from lysed cells/nuclei;
- **m***∈*ℝ^*G*^ is the bulk contamination profile, representing contamination introduced after pooling (i.e barcode-swapping and PCR chimeras);
- p ^k^ ∈ℝ ^G^ is the expected expression profile of cell-type *k*;
- *γ*_*i*_ *∈*{0, 1} ^*K*^ is the one-hot cell-type membership vector for barcode *i*, where 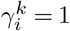 if barcode *i* is assigned to cell-type *k* and 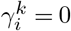 otherwise;
- *α*_*i*_ *∈*[0, 1] is the fraction of ambient RNA in barcode *i*;
- *β ∈* [0, 1] is the global bulk contamination fraction.

Each component profile **a, m, p**^*k*^ is normalized such that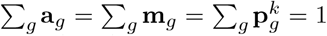The mixture weights *α*_*i*_,*β*, and *γ*_*i*_ thus represent true proportions and ensure that *χ*_*i*_ itself is a valid probability vector. Intuitively, this expression assumes that:

1. Barcode-specific contamination is governed by *α*_*i*_ (ambient RNA fraction);
2. Global contamination is governed by *β* (bulk library noise);
3. The biological cell signal is governed by an underlying cell-type profile indicated by *γ*_*i*_.

Together, these terms yield an interpretable and flexible mixture model.

### Modeling Assumptions

The model relies on the following assumptions:

- **Multinomial sampling:** Given the total UMI count *T*_*i*_, counts across genes follow a multinomial distribution with probabilities *χ*_**i**_. This ignores overdispersion but provides a tractable and empirically accurate approximation for large counts.
- **Linear mixing:** Counts from cell, ambient, and bulk sources combine additively prior to normalization. This reflects the physical mixture of molecules before UMI counting.
- **Fixed number of cell-types:** The dataset contains *K* latent cell-type expression profiles {**p**^*k*^}, which are shared across barcodes.
- **Barcode independence:** Each barcode is modeled independently given global parameters (**a, m**, {**p**^*k*^}, *β*).

## Initial Parameter Estimates

### Ambient Profile

In high-throughput single-cell assays, a large fraction of recovered barcodes are non-cellular, yet are nonetheless associated with small numbers of molecules originating from the ambient pool. Because the feature counts for these non-cellular barcodes originate solely from ambient contamination, these barcodes’ feature expression composition provides an unbiased estimate of the global ambient profile. For these technologies, we estimate the ambient distribution as the normalized average of counts across all barcodes identified as empty:

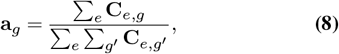

where *e* indexes the non-cellular barcodes.

Under the assumption that non-cellular barcodes sample independently from a shared ambient pool, this estimator corresponds to the maximum likelihood estimate of **a** under a multinomial sampling model, and is highly stable due to the large number of non-cellular barcodes typically observed.

### Bulk Contamination Profile

Across all technologies, a small fraction of global contamination can arise after pooling during library preparation and amplification. This contamination affects all barcodes approximately uniformly and can be modeled by a shared background profile **m**. Because this background reflects the aggregate expression composition of the entire experiment, we approximate **m** as the normalized mean expression profile across all droplets:

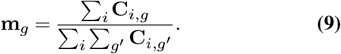

This estimator corresponds to the maximum likelihood estimate of the global background under a multinomial sampling model and provides a stable initialization for subsequent inference. In principle, an even more accurate estimate of **m** could be obtained from molecule counts prior to barcode collapsing, as bulk contamination is introduced during library preparation and amplification. However, such information is typically unavailable in standard scRNA-seq outputs, making the collapsed-count estimator a practical and robust approximation.

### Cell Typing

The number of cell-types *K* is specified or estimated from an initial clustering of the normalized count matrix. The cell-type assignment for each droplet is indicated by the one-hot cell membership vector *γ*_*i*_.

### Inference via Expectation–Maximization

We introduce a latent variable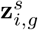 representing the number of UMI counts for barcode *i* and feature *g* that originated from source *s∈*{A, M, P_1_, …, *P*_*K*_}. The observed counts are constrained by:

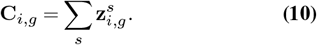

The likelihood is

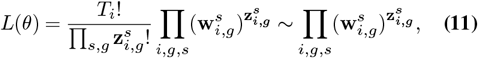

where*θ* = {{*α*_*i*_}, *β*, {**p**^*k*^}, **a, m**} and

- 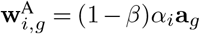
- 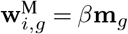
- 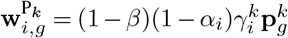

### E-step

In the expectation step, the latent counts are replaced by their expected values given the observed counts and current parameters:

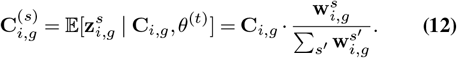

This can be interpreted as decomposing each observed count into fractional contributions from each source.

### M-step

The expected complete-data log-likelihood is

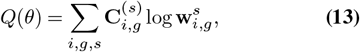

and parameters are updated by maximizing *Q* with respect to each group.

### Updating cell-type profiles

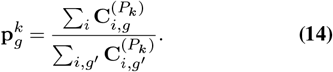

### Updating barcode contamination

We update the barcode-specific ambient fraction *α*_*i*_ from the expected ambient and cell-assigned counts:

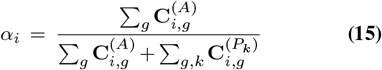

We do not update *α*_*i*_ for barcodes that were pre-classified as non-cellular: for such barcodes we keep *α*_*i*_ = 1 (i.e. 100% ambient, excluding the global bulk component).

### Updating the global bulk fraction

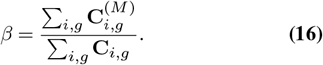

### Two-stage optimization and handling of extreme ambient fractions

CellSweep employs a two-stage optimization procedure designed to stabilize inference in the presence of barcodes that are poorly explained by their assigned cell-type profiles. Optimization begins with a burn-in phase, during which convergence is assessed exclusively via stabilization of the log-likelihood. This allows the algorithm to rapidly enter a stable basin of the likelihood surface before enforcing stricter parameter-based stopping criteria.During burn-in, per-barcode ambient fractions are capped at *α*_*i*_≤0.9. Barcodes that reach this cap are temporarily excluded from contributing to updates of the cell-type profiles *p*_*k*_, preventing nearly ambient barcodes from biasing early profile estimates. Only these capped barcodes are permitted to update their cell-type assignments during burn-in, using a hard maximum-likelihood criterion over existing profiles. After convergence of the log-likelihood, the ambient cap is removed, cell-type assignments are fixed, and final optimization proceeds until convergence of the model parameters. In practice, this procedure substantially reduces the prevalence of extreme *α*_*i*_→ 1 solutions while preserving global clustering structure and stabilizing estimation of the cell-type profiles.

### Repulsion-modified update for cell-type profiles

During the burn-in phase, CellSweep modifies the standard EM update for cell-type expression profiles to discourage solutions in which inferred cell-type profiles align closely with the ambient profile. Rather than maximizing the expected complete-data log-likelihood alone, the cell-type profiles are updated by solving a penalized optimization problem that explicitly disfavors similarity to the ambient profile. For a given cell-type profile *p∈* Δ^*G*−1^, the burn-in update is defined as the solution to

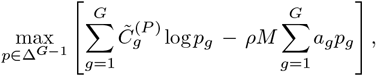

where 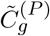denotes the expected number of counts assigned to the cell-type component for feature *g*, 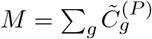 the total expected mass of the profile, *a* is the fixed ambient expression profile, and *ρ >* 0 controls the strength of the repulsion. This modified objective biases early updates away from explaining background signal as biological variation. After the burn-in phase, the repulsion term is disabled and standard EM updates are recovered exactly, ensuring that final estimates correspond to a stationary point of the original likelihood.

### Inferring the Ambient Profile without Non-Cellular Barcodes

In situations with few or no non-celluar barcodes, the ambient pool can instead be viewed as a mixture of cell-type profiles. Conceptually, ambient contamination arises from lysed or damaged cells, so the global ambient profile should lie within the convex hull of the cell-type expression profiles {*p*^*k*^}.

We therefore parameterize the ambient profile as a mixture of the *K* cell-type expression distributions:

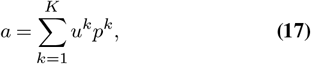

such that *u*^*k*^ satisfies 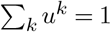 and *u*^*k*^ > 0.We initialize these mixture weights using the initial soft membership assignments 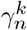Specifically, we set

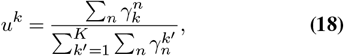

which corresponds to the estimated relative abundance of each cell type in the dataset. This choice ensures that the initial ambient profile resembles the global expression distribution implied by the initial clustering, while remaining flexible enough to adapt during subsequent updates.

Allowing the ambient profile to be updated renders the two-step optimization strategy used in the default model ineffective. In the default setting, the ambient profile is fixed and therefore acts as a stable anchor against which signal can be distinguished from noise. When the ambient profile is allowed to change, the inferred structure of the noise coevolves with the cell-type profiles, so it no longer provides a consistent reference direction for separating signal from background. In this regime, repulsion and cell-type reassignment become unstable because similarity to ambient is itself moving during training. For this reason, the alternative model omits both the repulsion term and cell-type reassignment.

### Updating the ambient mixture weights

When non-cellular barcodes are unavailable, the ambient mixture weights {*u*^*k*^}are refined using a nested expectation–maximization procedure. Let *A*_*g*_ denote the expected number of counts assigned to the ambient component for feature *g*, aggregated across all barcodes. Given the current estimate of the ambient profile ***a*** and the cell-type profiles {***p***^*k*^}, we compute the posterior responsibility of each cell type for generating ambient counts at each feature,

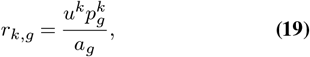

These responsibilities are then used to update the mixture weights,

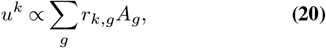

followed by normalization to enforce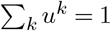. The ambient profile is subsequently recomputed as

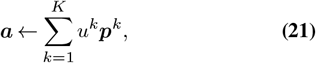

.This inner update is repeated for three iterations to refine the ambient mixture weights without fully optimizing them at each outer EM step. This approach provides a stable compromise between flexibility and computational efficiency, allowing the ambient profile to adapt while avoiding oscillatory behavior.

### Recovering Denoised Counts

After convergence, the cleaned data matrix is returned by subtracting the expected noise counts from the observed counts:

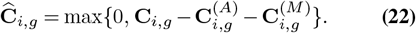

When integer counts are required, stochastic rounding is used to preserve unbiasedness and per-barcode totals.

## Notes

### Competing Interest Statement

The authors have declared no competing interest.

