## Supplementary Figures for "Single-Cell Genomics Decontamination with CellSweep"

### Supplementary Figures
### Single-Cell Genomics Decontamination with CellSweep

Maya Caskey[1,*], Joseph Rich[1,*], Ryan Weber[2], Ali Mortazavi[2], Lior Pachter[1], and Ingileif Hallgrimsdottir[1,†]

[1]Division of Biology and Biological Engineering, California Institute of Technology, Pasadena, California, USA, 91125
[2]Department of Developmental and Cell Biology, University of California, Irvine, Irvine, California, USA, 92697
[*]These authors contributed equally

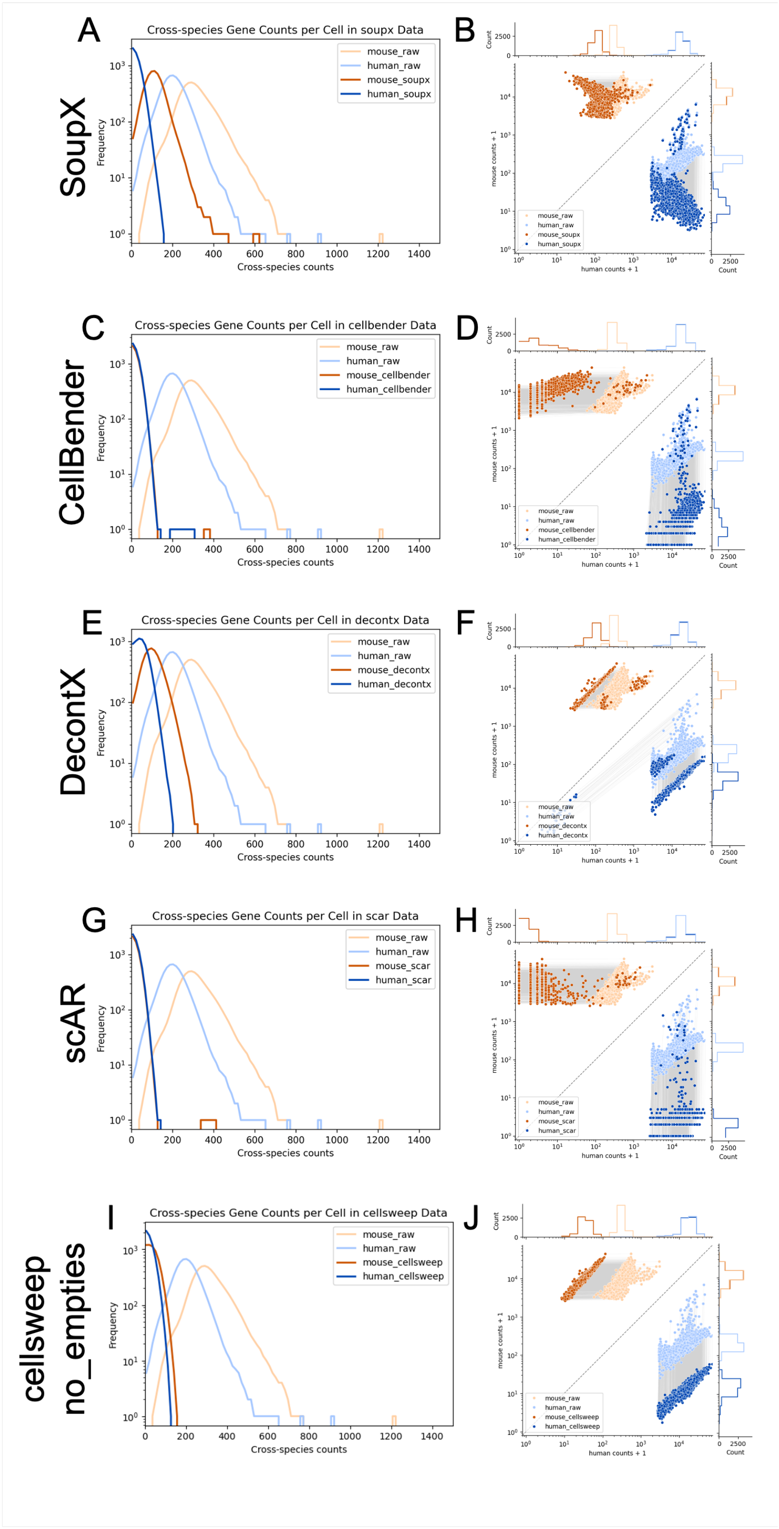


**Fig S1.** Analysis of the human-mouse mixture dataset with additional tools. (A) Histogram of total cross-species counts across all genes per cell after processing with SoupX. (B) Joint scatterplot of mouse vs. human counts after processing with SoupX. (C-D) Same as (A-B) but with CellBender. (E-F) Same as (A-B) but with DecontX. (G-H) Same as (A-B) but with scAR. (I-J) Same as (A-B) but with the alternative CellSweep model after empty droplets are removed. Light orange = mouse cells, raw; light blue = human cells, raw; dark orange = mouse cells, processed; dark blue = human cells, processed.

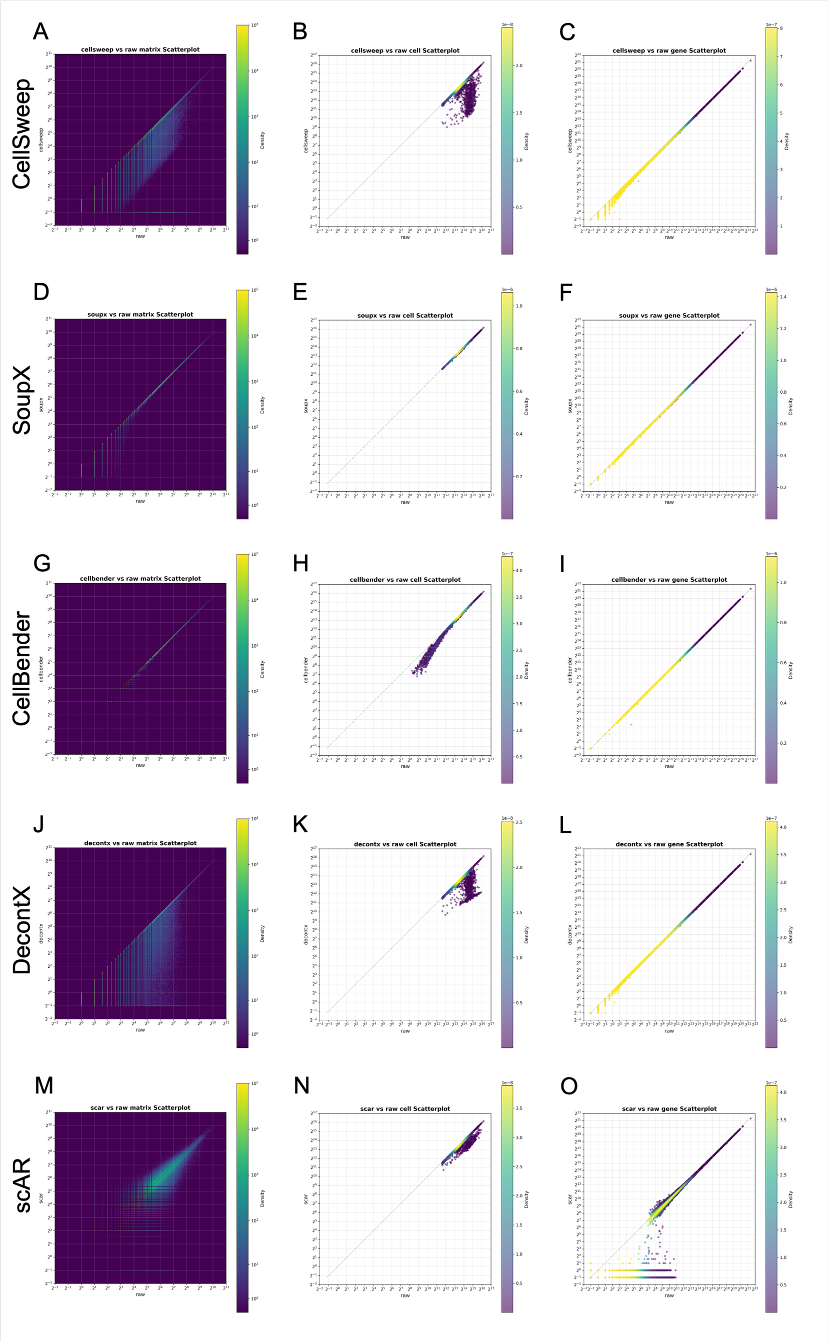


**Fig S2.** Additional scatterplots for the analysis of the human-mouse mixture dataset. (A-C) Scatterplot of matrix counts (A), total cell counts (B), and total gene counts (C) after vs. before processing with CellSweep. (D-F) Same as (A-C) but with SoupX. (G-I) Same as (A-C) but with CellBender. (J-L) Same as (A-C) but with DecontX. (M-O) Same as (A-C) but with scAR.

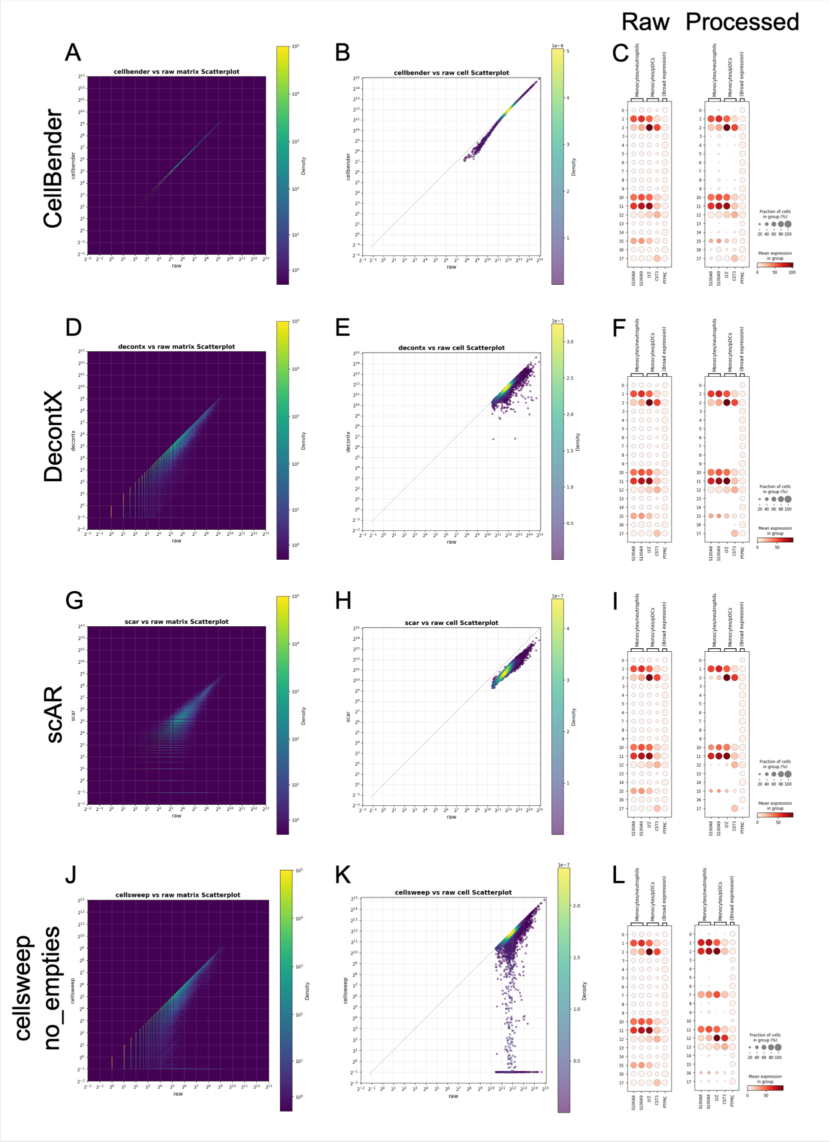


**Fig S3.** Analysis of the PBMC 8k dataset with additional tools. (A) Scatterplot of matrix values after vs. before processing with CellBender. (B) Scatterplot of total cell counts after vs. before processing with CellBender. (C) Dotplots of markers from monocytes/neutrophils, monocytes/pDCs, and broad expression in raw (left) and processed (right) data with CellBender. (D-F) Same as (A-C) but with DecontX. (G-I) Same as (A-C) but with scAR. (J-L) Same as (A-C) but with the alternative CellSweep model after empty droplets are removed

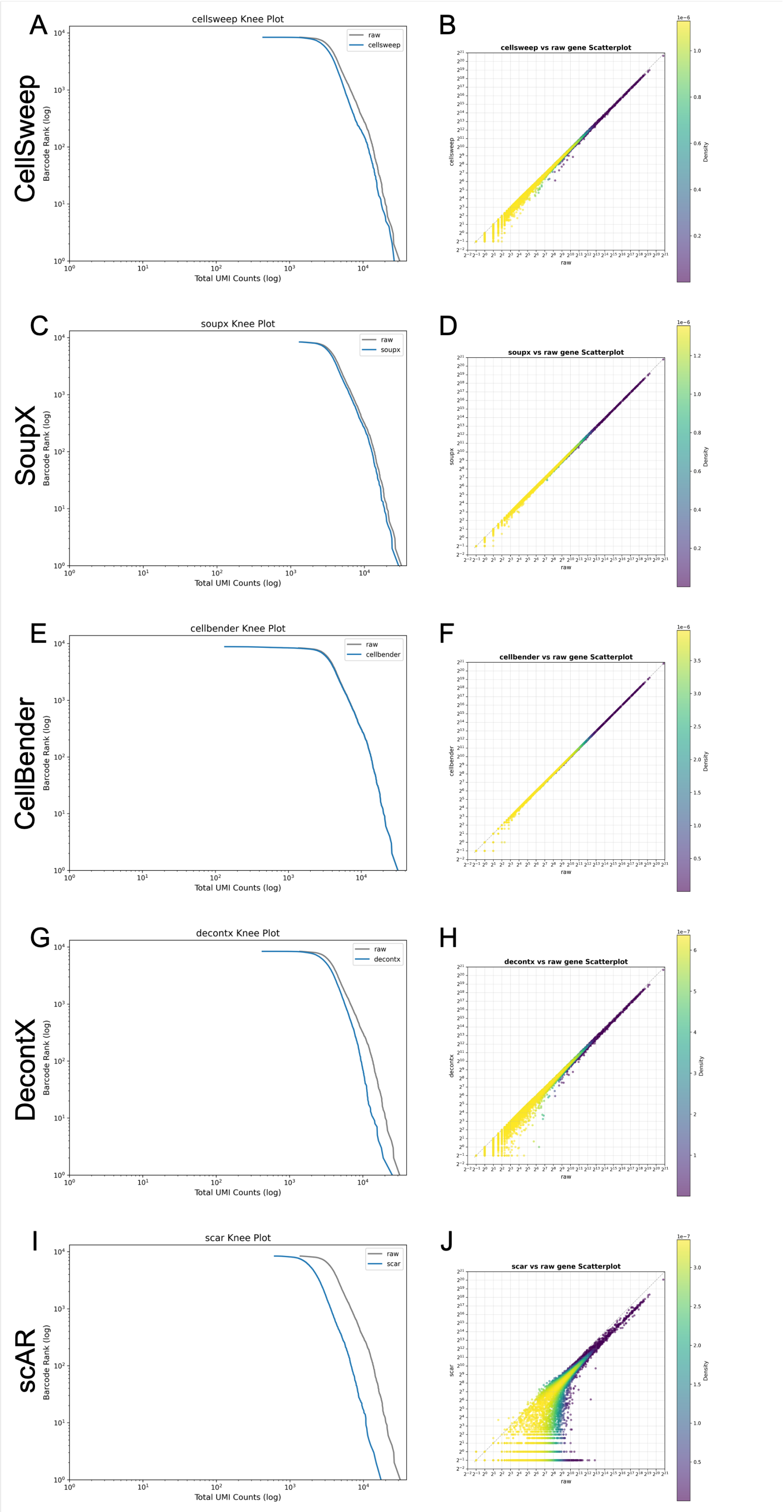


**Fig S4.** Knee plots and gene scatterplots for the analysis of the PBMC 8k dataset. (A) Knee plots of cell-containing droplets before and after processing with CellSweep. (B) Scatterplot of total gene counts after vs. before processing with CellSweep. (C-D) Same as (A-B) but with SoupX. (E-F) Same as (A-B) but with CellBender. (G-H) Same as (A-B) but with DecontX. (I-J) Same as (A-B) but with scAR.

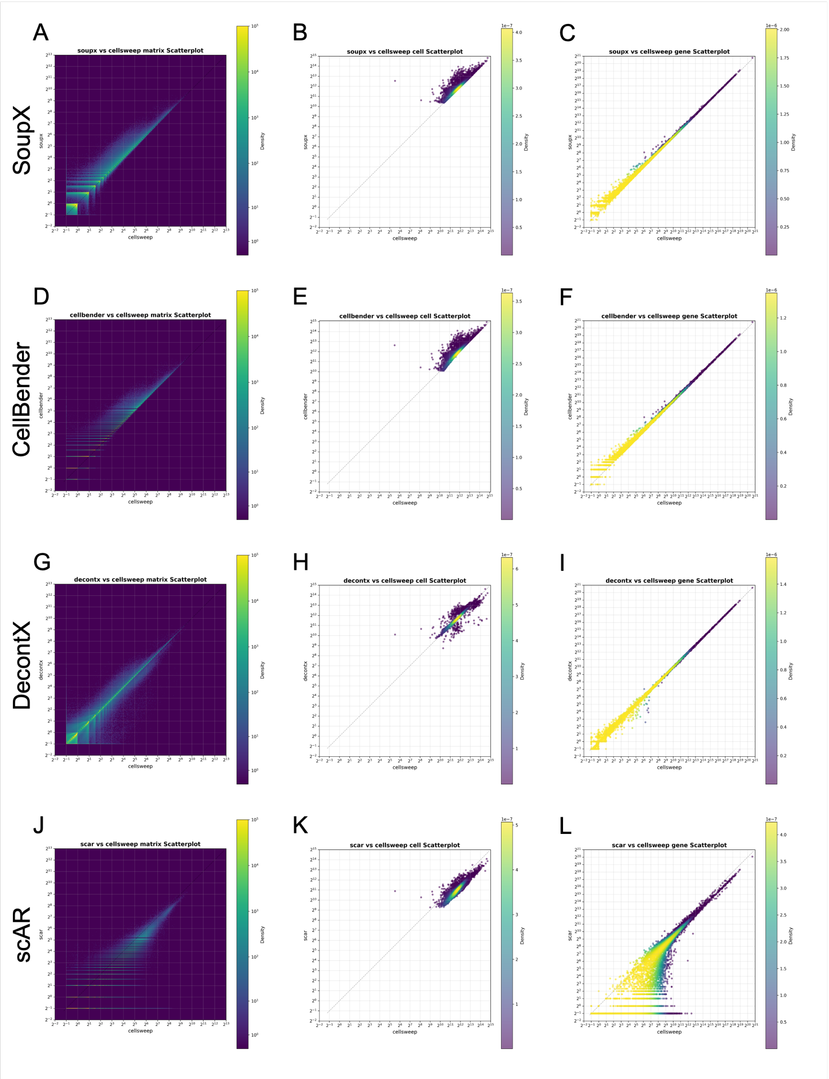


**Fig S5.** Comparison of CellSweep to other tools on the PBMC 8k dataset. (A-C) Scatterplot of matrix counts (A), total cell counts (B), and total gene counts (C) after processing with SoupX vs. CellSweep. (D-F) Same as (A-C) but with CellBender. (G-I) Same as (A-C) but with DecontX. (J-L) Same as (A-C) but with scAR.

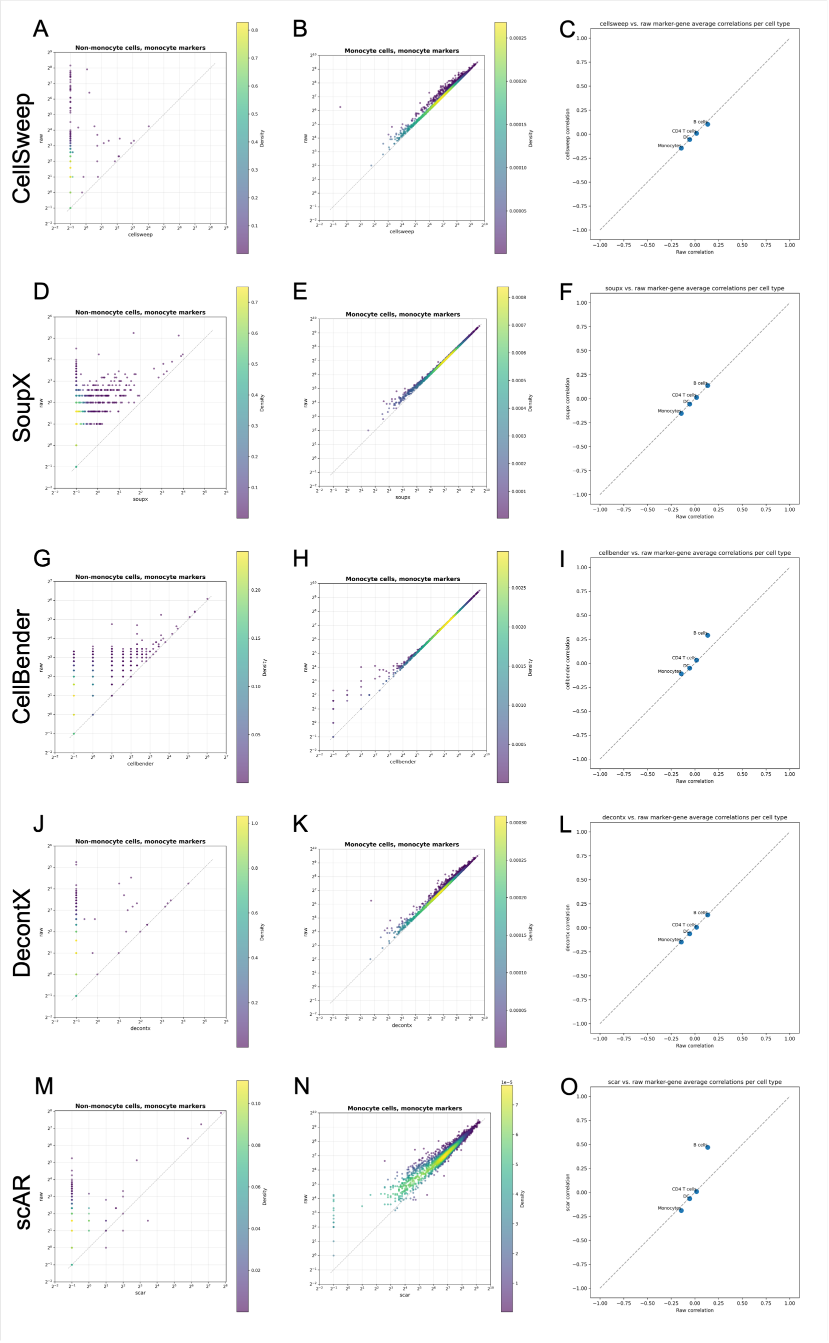


**Fig S6.** Additional marker gene scatterplots for analysis of the PBMC 8K dataset. (A) Scatterplot of total monocyte marker gene expression in non-monocyte cells after vs. before processing with CellSweep. (B) Scatterplot of total monocyte marker gene expression in monocyte cells after versus before processing with self sleep. (C) Mean gene-gene Pearson correlation for marker genes of a given cell type after vs. before processing with CellSweep. (D-F) Same as (A-C) but with SoupX. (G-I) Same as (A-C) but with CellBender. (J-L) Same as (A-C) but with DecontX. (M-O) Same as (A-C) but with scAR.

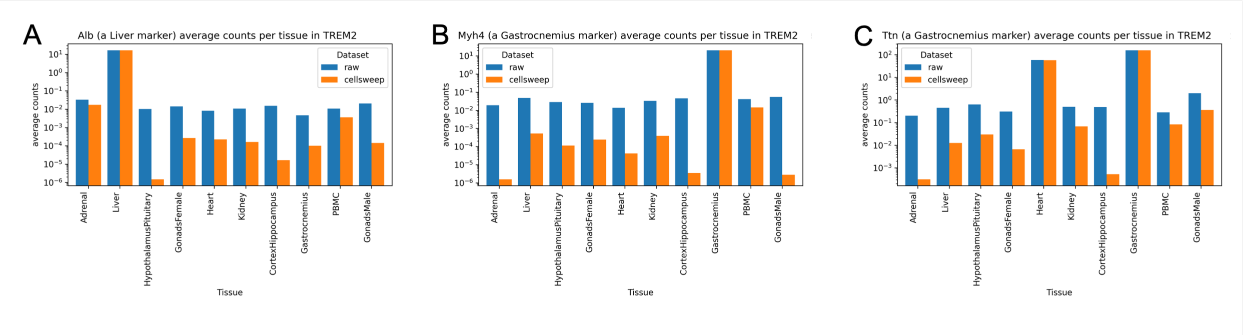


**Fig S7.** Cellsweep reduces non-marker gene counts in 8 cubed Trem2 data. Bar plots depict expression across tissues of the liver marker Alb (A) and the muscle markers Myh4 (B) and Ttn (C).

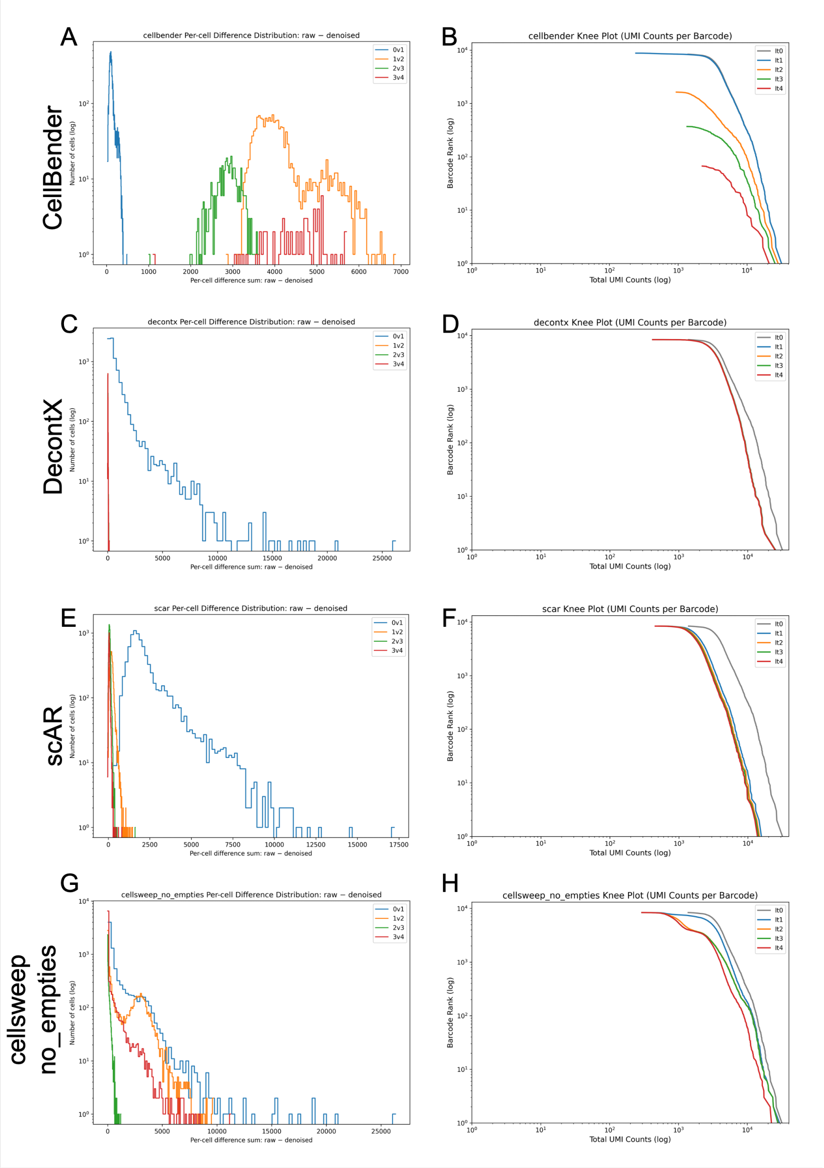


**Fig S8.** Idempotency analysis with additional tools. (A) Histogram of per-cell count differences after processing with CellBender. Blue = between iteration 0 (raw) and 1; orange = between iteration 1 and 2; green = between iteration 2 and 3; red = between iteration 3 and 4. (D) Knee plots after processing with CellBender. Gray = iteration 0 (raw). Blue = iteration 1; orange = iteration 2; green = iteration 3; red = iteration 4. (C-D) Same as (A-B) but with DecontX. (E-F) Same as (C-D) but with DecontX. (G-H) Same as (A-B) but with the alternative CellSweep model after empty droplets are removed.

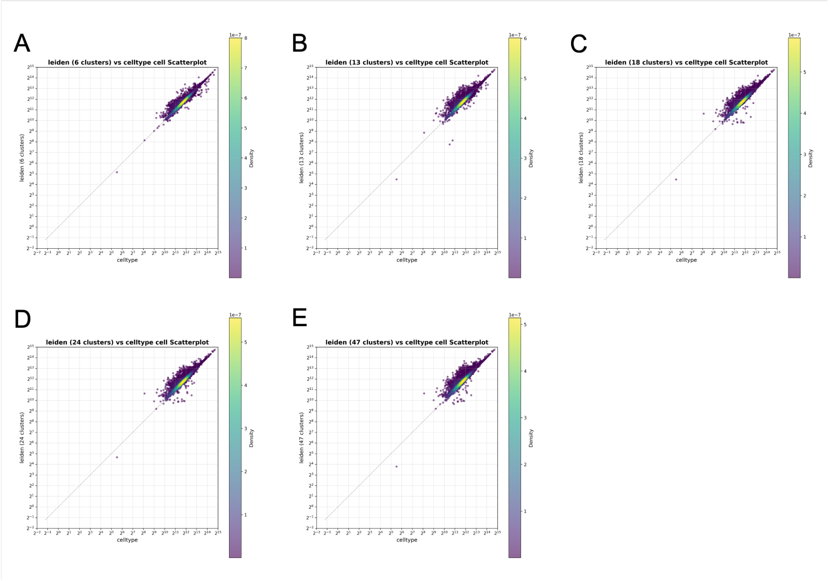


**Fig S9.** Celltypes determined by Leiden clustering vs. CellTypist. Repeated with various Leiden resolutions resulting in 6 clusters (A), 13 clusters (B), 18 clusters (C), 24 clusters (D), and 47 clusters (E). Analysis performed on the PBMC 8k dataset.

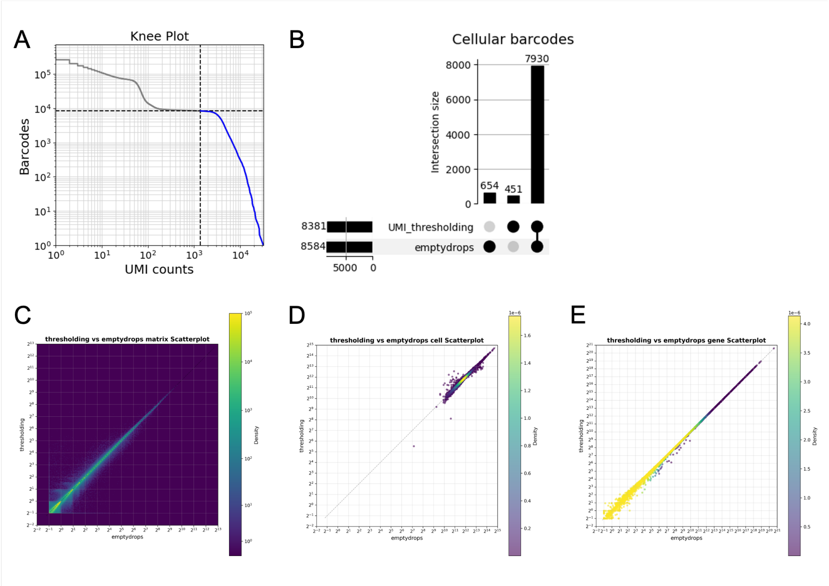


**Fig S10.** Sensitivity to empty barcode identification method. (A) Knee plot. (B) UpSet plot comparing cell-containing barcodes between UMI thresholding with 8,381 cells (expected cells on the 10x website) and EmptyDrops. (C) Scatterplot of matrix (C) counts (C), total cell counts (D), and total gene counts (E) after running CellSweep with empty barcodes identified by thresholding vs. EmptyDrops. Analysis performed on the PBMC 8k dataset.

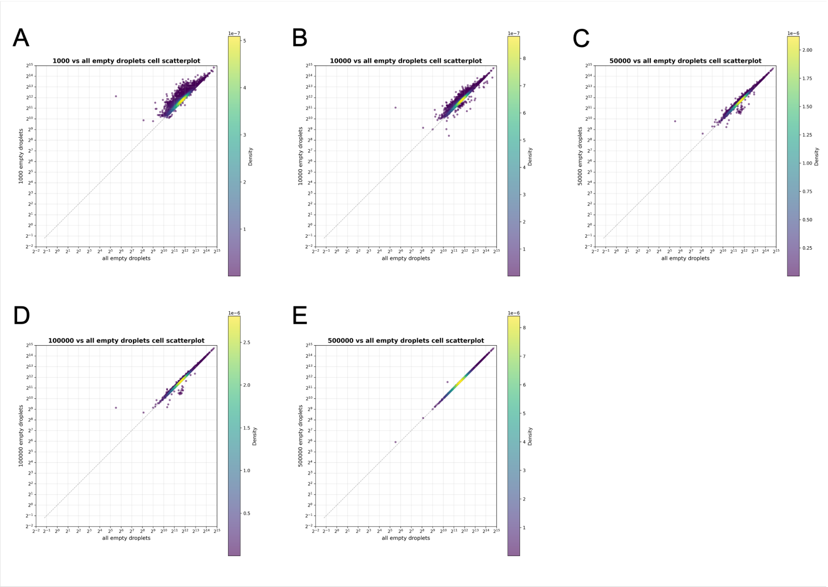


**Fig S11.** Sensitivity to number of empty barcodes. Scatterplots of total counts per cell with subsampled empty barcodes vs. all (728,899) empty barcodes. Repeated with 1,000 (A), 10,000 (B), 50,000 (C), 100,000 (D), and 500,000 (E) empty barcodes. Analysis performed on the PBMC 8k dataset.

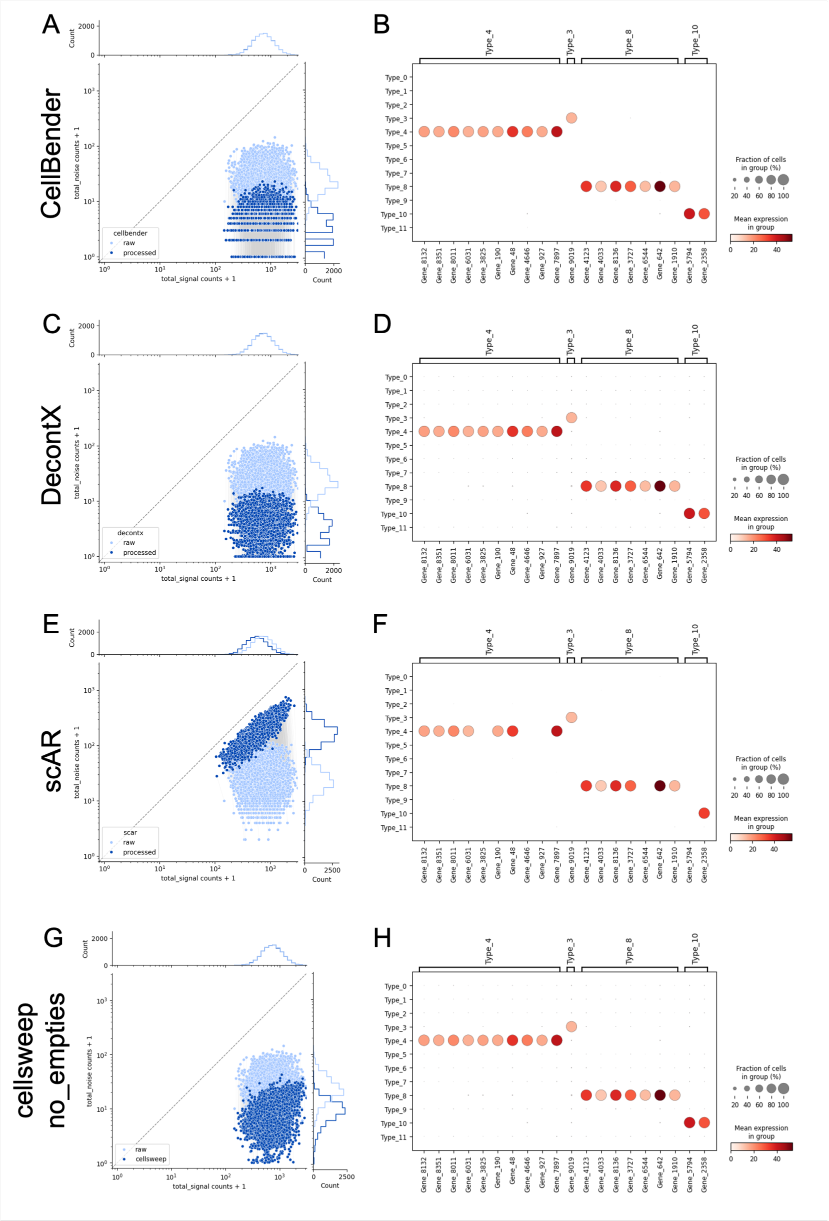


**Fig S12.** Analysis of the simulated dataset with additional tools. (A) Joint scatterplot of noise and signal total marker counts per cell after processing with CellBender. Light blue = raw; dark blue = CellBender. (B) Dotplot of processed data with CellBender. (C-D) Same as (A-B) but with DecontX. (E-F) Same as (A-B) but with scAR. (G-H) Same as (A-B) but with the alternative CellSweep model after removal of non-cellular barcodes.

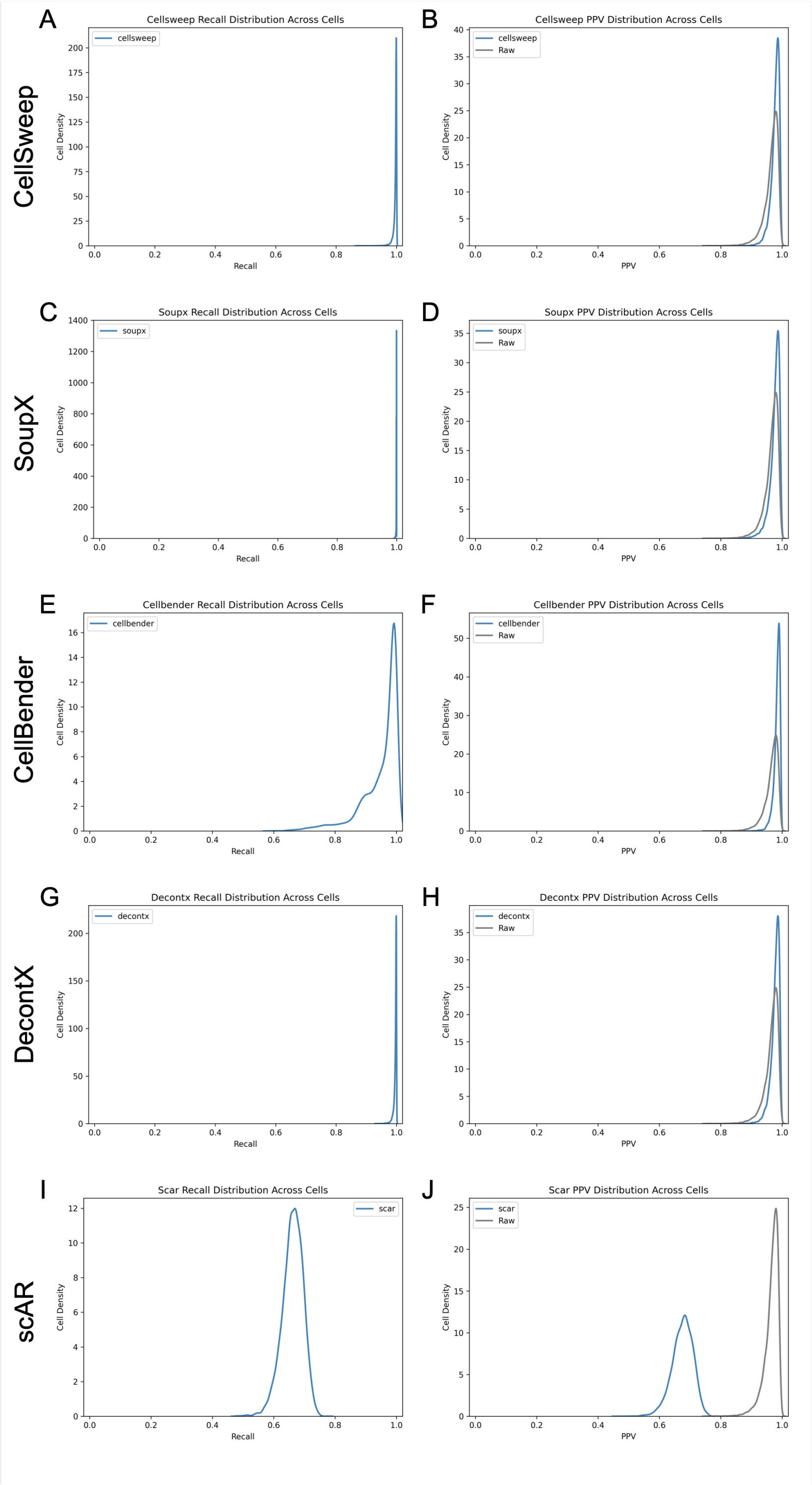


**Fig S13.** Additional plots for the simulated dataset. (A-B) Smooth density curves for sensitivity (A) and positive predictive value (B) per cell for raw data and data processed with CellSweep. (C-D) Same as (A-B) but with SoupX. (E-F) Same as (A-B) but with CellBender. (G-H) Same as (A-B) but with DecontX. (I-J) Same as (A-B) but with scAR.

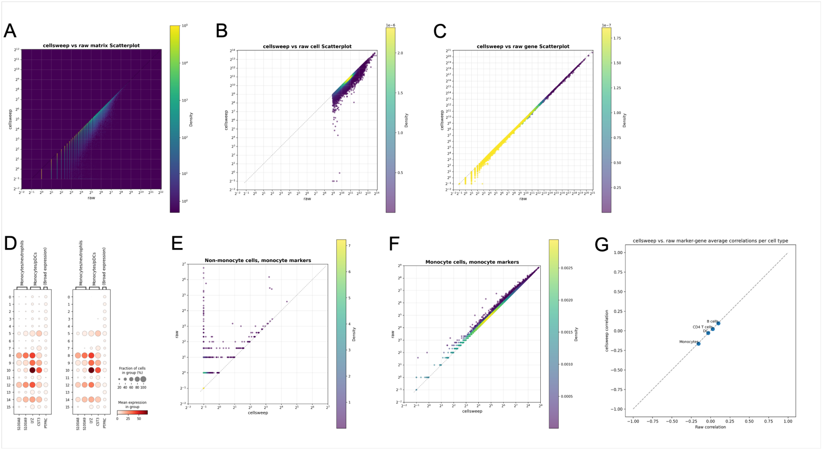


**Fig S14.** Analysis of human PBMC 33k scRNA-seq dataset with CellSweep. (A-C) Scatterplot of matrix counts (A), total cell counts (B), and total gene counts (C) after vs. before processing with CellSweep. (D) Dotplots of markers from monocytes/neutrophils, monocytes/pDCs, and broad expression in raw (left) and processed (right) data with CellSweep. (E) Scatterplot of total monocyte marker gene expression in non-monocyte cells after vs. before processing with CellSweep. (F) Scatterplot of total monocyte marker gene expression in monocyte cells after versus before processing with self sleep. (G) Mean gene-gene Pearson correlation for marker genes of a given cell type after vs. before processing with CellSweep.

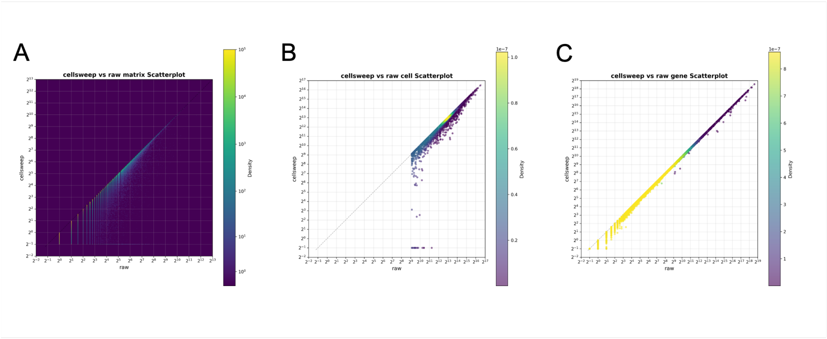


**Fig S15.** Analysis of mouse PBMC scRNA-seq dataset with CellSweep. (A-C) Scatterplot of matrix counts (A), total cell counts (B), and total gene counts (C) after vs. before processing with CellSweep.

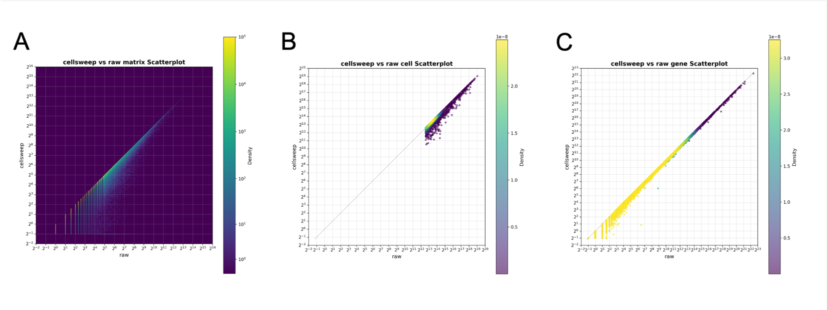


**Fig S16.** Analysis of human melanoma scRNA-seq dataset with CellSweep. (A-C) Scatterplot of matrix counts (A), total cell counts (B), and total gene counts (C) after vs. before processing with CellSweep.

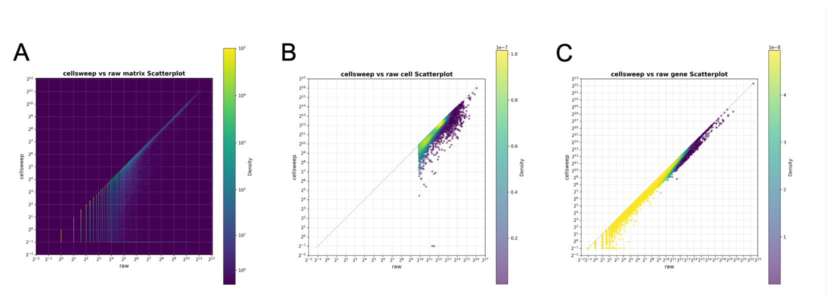


**Fig S17.** Analysis of human kidney single-nucleus RNA-seq dataset with CellSweep. (A-C) Scatterplot of matrix counts (A), total cell counts (B), and total gene counts (C) after vs. before processing with CellSweep.
