## Supplementary Methods for "Single-Cell Genomics Decontamination with CellSweep"

### The CellSweep Model

#### Model Notation.

##### Observed quantities

- $\mathbf{C}_i \in \mathbb{Z}_+^G$: observed count vector for barcode $i$
- $T_i \in \mathbb{Z}_+$: total observed counts for barcode $i$

##### Fixed input

- $\gamma_i \in \{0,1\}^K$: cell-type assignment for barcode $i$, where $\gamma_i^k = 1$ if barcode $i$ is assigned to cell-type $k$ and $\gamma_i^k = 0$ otherwise

##### Estimated parameters

- $\chi_i \in \mathbb{R}_+^G$: expected count profile for barcode $i$, normalized such that $\sum_g \chi_{i,g} = 1$
- $\mathbf{a} \in \mathbb{R}_+^G$: ambient contamination profile, normalized such that $\sum_g a_g = 1$
- $\mathbf{m} \in \mathbb{R}_+^G$: bulk contamination profile, normalized such that $\sum_g m_g = 1$
- $\mathbf{p}^k \in \mathbb{R}_+^G$: expected expression profile of cell type $k$, normalized such that $\sum_g p_g^k = 1$
- $\alpha_i \in [0,1]$: barcode-specific ambient contamination fraction
- $\beta \in [0,1]$: global bulk contamination fraction

**Statistical Model.** We model the counts for a barcode $i$ as

$$\mathbf{C}_i \sim \text{Multinomial}(T_i, \chi_i) \tag{1}$$

where, $T_i$ is the total counts in barcode $i$, and

$$\chi_i = (1-\beta)\big[\alpha_i \mathbf{a} + (1-\alpha_i)\sum_{k=1}^{K} \gamma_i^k \mathbf{p^k}\big] + \beta \mathbf{m}. \tag{2}$$

***Parameter Initialization.***

- $\alpha_i = 0.9$ for all cell-containing barcodes. For non-cellular barcodes, $\alpha_i$ is fixed at 1.
- $\beta = 0.1$. This value is initialized above the expected bulk contamination fraction to avoid convergence to a degenerate zero solution during training.
- $\gamma_i^k = 1$ if barcode $i$ is assigned to cell-type $k$ and $\gamma_i^k = 0$ otherwise.
- $$\mathbf{a}_g = \frac{\sum_e \mathbf{C}_{e,g} + \frac{50}{G}}{\sum_{g'}(\sum_e \mathbf{C}_{e,g'} + \frac{50}{G})}, \tag{3}$$
  where $e$ indexes barcodes classified as non-cellular. The term $\frac{50}{G}$ acts as a small uniform Dirichlet prior for numerical stability.
- $$\mathbf{m}_g = \frac{\sum_i \mathbf{C}_{i,g} + \frac{10}{G}}{\sum_{g'}(\sum_i \mathbf{C}_{i,g'} + \frac{10}{G})}. \tag{4}$$
  The term, $\frac{10}{G}$ acts as a small uniform Dirichlet prior for numerical stability.
- For cell-type $k$:
  $$\mathbf{p}_g^k = \frac{\sum_i \gamma_i^k \mathbf{C}_{i,g} + \frac{50}{G}}{\sum_{g'}(\sum_i \gamma_i^k \mathbf{C}_{i,g'} + \frac{50}{G})}. \tag{5}$$
  The term, $\frac{50}{G}$ acts as a small uniform Dirichlet prior for numerical stability.

**Defining Cell-Types.** Cell-type assignments are required input. In this paper, unless otherwise specified, CellTypist is used.

**Likelihood terminated burn-in phase.** Optimization is divided into an initial burn-in phase and a final refinement phase using a likelihood-based criterion. At each iteration, we monitor the absolute change in the average per-barcode log-likelihood. An adaptive tolerance is initialized after the second iteration as a fixed fraction of the initial likelihood change. To prevent the tolerance from collapsing to unrealistically small values as the likelihood scale varies across datasets, we additionally enforce a minimum absolute threshold proportional to the magnitude of the likelihood via a relative tolerance. The effective tolerance at iteration $t$ is taken as the maximum of these two quantities.

Formally, let $\ell^{(t)}$ denote the average per-barcode log-likelihood at iteration $t$. The adaptive tolerance is defined as

$$\tau^{(t)} = \max\Big(10^{-3}\left|\ell^{(2)} - \ell^{(1)}\right|,\, 10^{-6}\max\Big(|\ell^{(t-1)}|, 1\Big)\Big),$$

and the burn-in phase is terminated once

$$\left|\ell^{(t)} - \ell^{(t-1)}\right| < \tau^{(t)}.$$

The relative threshold $10^{-6}$ reflects the effective precision limit of 32-bit floating point arithmetic for accumulated log-likelihood values and prevents convergence checks from being driven by numerical noise rather than meaningful parameter updates.

Empirically, the log-likelihood stabilizes rapidly and consistently across datasets, typically well before individual parameters. We therefore use likelihood stabilization exclusively to terminate the burn-in phase, signaling that the optimization has entered a stable basin. Final convergence is assessed separately using stricter criteria based on changes in $p_k$ and the effective contamination fraction, ensuring accurate parameter estimates without unnecessarily prolonging burn-in.

**Expectation–Maximization.** Let $C_{i,g} \in \mathbb{N}$ denote the observed UMI count for barcode $i \in \{1,\ldots,N\}$ and feature $g \in \{1,\ldots,G\}$. We introduce latent allocation variables

$$z^s_{i,g} \in \mathbb{N}, \qquad s \in \mathcal{S} := \{A, M, P_1, \ldots, P_K\},$$

representing the number of counts of feature $g$ in barcode $i$ originating from source $s$. These variables satisfy the constraint

$$\sum_{s\in\mathcal{S}} z^s_{i,g} = C_{i,g}. \tag{6}$$

***Complete-data likelihood.*** The complete-data likelihood factorizes as a product of multinomial distributions when conditioned on the mixture weights $\{w^s_{i,g}\}_{s\in\mathcal{S}}$:

$$p(\{z^s_{i,g}\} \mid \theta) = \prod_{i,g} \frac{C_{i,g}!}{\prod_{s\in\mathcal{S}} z^s_{i,g}!} \prod_{s\in\mathcal{S}} \left(w^s_{i,g}\right)^{z^s_{i,g}}, \tag{7}$$

where the parameters are

$$\theta = \Big(\{\alpha_i\}_{i=1}^N, \beta, \{\mathbf{p}^k\}_{k=1}^K\Big)$$

and the component weights are defined by

$$w^A_{i,g} = (1-\beta)\alpha_i a_g, \tag{8}$$

$$w^M_{i,g} = \beta m_g, \tag{9}$$

$$w^{P_k}_{i,g} = (1-\beta)(1-\alpha_i)\gamma^k_i p^k_g. \tag{10}$$

By construction,

$$\sum_{s\in\mathcal{S}} w^s_{i,g} = 1 \quad \forall i, g.$$

Ignoring terms independent of $\theta$, the complete-data log-likelihood is

$$\ell_c(\theta) = \sum_{i,g,s} z^s_{i,g} \log w^s_{i,g}. \tag{11}$$

***E-step.*** At iteration $t$, the E-step computes the conditional expectation of the latent variables under the current parameter values $\theta^{(t)}$:

$$C^{(s)}_{i,g} := \mathbb{E}\Big[z^s_{i,g} \mid C_{i,g}, \theta^{(t)}\Big].$$

Since $(z^s_{i,g})_{s\in\mathcal{S}} \mid C_{i,g}$ follows a multinomial distribution with probabilities proportional to $w^s_{i,g}$, we obtain

$$C^{(s)}_{i,g} = C_{i,g} \frac{w^s_{i,g}}{\sum_{s'\in\mathcal{S}} w^{s'}_{i,g}}. \tag{12}$$

***M-step.*** The M-step maximizes the expected complete-data log-likelihood

$$Q(\theta \mid \theta^{(t)}) := \mathbb{E}_{\theta^{(t)}}[\ell_c(\theta)] = \sum_{i,g,s} C^{(s)}_{i,g} \log w^s_{i,g} \tag{13}$$

with respect to $\theta$, subject to the normalization constraints on $\mathbf{p}^k$ and $\mathbf{a}$.

**Update of cell-type profiles $\mathbf{p}^k$.** For fixed $k$, the relevant terms of $Q$ are

$$Q(\mathbf{p}^k) = \sum_{i,g} C^{(P_k)}_{i,g} \log p^k_g \quad \text{subject to} \quad \sum_g p^k_g = 1.$$

Introducing a Lagrange multiplier $\lambda_k$ and differentiating yields

$$\frac{\partial}{\partial p^k_g}\left(\sum_{i,g} C^{(P_k)}_{i,g} \log p^k_g + \lambda_k\left(1 - \sum_g p^k_g\right)\right) = \frac{\sum_i C^{(P_k)}_{i,g}}{p^k_g} - \lambda_k = 0.$$

Solving for $p^k_g$ and enforcing normalization gives the maximum likelihood update

$$p^k_g = \frac{\sum_i C^{(P_k)}_{i,g}}{\sum_{i,g'} C^{(P_k)}_{i,g'}}. \tag{14}$$

In practice, the likelihood admits near-degenerate solutions in which one or more inferred cell-type profiles drift toward the ambient expression profile. In this regime, background counts can be explained by the cell-type component rather than the ambient component, often driving the estimated ambient fractions toward implausibly small values. To discourage this failure mode, we introduce a repulsion modification to the cell-type profile update during an initial burn-in stage. After burn-in, the repulsion is disabled and inference proceeds under the original likelihood.

***Burn-in Repulsion Modified Update.*** We consider the M-step update of a single cell-type profile $p \in \Delta^{G-1}$, where $\Delta^{G-1}$ denotes the probability simplex. Let $\tilde{C}_g^{(P)}$ denote the expected number of counts assigned to gene $g$ from the cell-profile component by the E-step, and let

$$M = \sum_{g=1}^{G} \tilde{C}_g^{(P)}$$

be the corresponding total expected count mass.

**Penalized objective.** To discourage alignment between the cell-type profile and the ambient profile $a \in \Delta^{G-1}$, a natural penalized M-step objective is

$$\max_{p\in\Delta^{G-1}} \left[ \sum_{g=1}^{G} \tilde{C}_g^{(P)} \log p_g - \rho M \sum_{g=1}^{G} a_g p_g \right], \tag{15}$$

where $\rho > 0$ controls the strength of the repulsion. The objective Eq. (15) does not admit a simple closed-form maximizer and, if optimized exactly, can induce large and unstable updates early in EM when responsibilities are unreliable.

**Exact EM anchor point.** Rather than optimizing Eq. (15) directly, we first compute the exact EM M-step solution for the likelihood term alone:

$$p^{(0)} = \arg\max_{p\in\Delta^{G-1}} \sum_{g=1}^{G} \tilde{C}_g^{(P)} \log p_g = \frac{\tilde{C}^{(P)}}{M}. \tag{16}$$

This solution maximizes the expected complete-data log-likelihood given the current E-step responsibilities and serves as a stable reference point.

**Trust-region correction.** We incorporate the repulsion term as a local correction around $p^{(0)}$, rather than as a global re-optimization. Specifically, we restrict the influence of the penalty by imposing a quadratic trust region centered at $p^{(0)}$, leading to the following correction subproblem:

$$p^{\text{new}} = \arg\max_{p\in\Delta^{G-1}} \left[ -\rho M \sum_{g=1}^{G} a_g p_g - \frac{M}{2} \|p - p^{(0)}\|_2^2 \right]. \tag{17}$$

The trust-region penalty is scaled by the expected cluster mass $M$ to match the scale of the likelihood term, yielding depth-invariant updates and an effective proximal step size that decreases with increasing statistical support.

**Closed-form solution.** Dividing Eq. (17) by $M > 0$ and completing the square yields

$$p^{\text{new}} = \arg\min_{p\in\Delta^{G-1}} \left\| p - \left(p^{(0)} - \rho a\right) \right\|_2^2,$$

which is the Euclidean projection onto the simplex. Therefore,

$$p^{\text{new}} = \Pi_{\Delta^{G-1}} \left( p^{(0)} - \rho a \right). \tag{18}$$

**Count-space form.** Writing $q = Mp$, the update Eq. (18) is equivalently expressed in count space as

$$q^{\text{new}} = \Pi_{\{q\geq 0,\, \sum q = M\}} \left( \tilde{C}^{(P)} - (\rho M) a \right), \qquad p^{\text{new}} = \frac{q^{\text{new}}}{M}.$$

In practice, we apply a symmetric Dirichlet-regularized update for numerical stability by replacing $\tilde{C}^{(P)}$ with $\tilde{C}^{(P)} + \lambda$, where $\lambda = 50/G$ corresponds to a uniform Dirichlet prior with total mass 50.

To avoid numerical artifacts arising from repeated clipping at the boundary of the simplex, we additionally cap the repulsion-induced reduction for each gene. Specifically, the subtraction applied to gene $g$ is limited to a fixed fraction $\eta$ of its current count mass, with $\eta = 0.2$. This constraint prevents any single gene's expected count from being reduced by more than 20% in a single iteration and serves purely as a numerical safeguard.

**Interpretation.** This update can be viewed as an exact EM M-step for the likelihood followed by a single proximal (trust-region) correction for the repulsion penalty. The quadratic term does not approximate the likelihood; rather, it constrains the magnitude of the penalty-induced update, ensuring stable behavior during early iterations. After the burn-in phase, the repulsion term is disabled and standard EM updates are recovered.

**Update of barcode-specific ambient fractions $\alpha_i$.** Collecting terms involving $\alpha_i$ yields

$$Q(\alpha_i) = \sum_{g} C_{i,g}^{(A)} \log \alpha_i + \sum_{g,k} C_{i,g}^{(P_k)} \log(1-\alpha_i) + \text{const.}$$

Taking the derivative and setting it to zero:

$$\frac{\partial Q}{\partial \alpha_i} = \frac{\sum_g C_{i,g}^{(A)}}{\alpha_i} - \frac{\sum_{g,k} C_{i,g}^{(P_k)}}{1-\alpha_i} = 0,$$

which yields the closed-form solution

$$\alpha_i = \frac{\sum_g C_{i,g}^{(A)}}{\sum_g C_{i,g}^{(A)} + \sum_{g,k} C_{i,g}^{(P_k)}}. \tag{19}$$

For barcodes pre-classified as non-cellular, $\alpha_i$ is fixed to 1.

**Update of the global bulk fraction $\beta$.** The terms involving $\beta$ are

$$Q(\beta) = \sum_{i,g} C_{i,g}^{(M)} \log \beta + \sum_{i,g} \left( C_{i,g}^{(A)} + \sum_{k} C_{i,g}^{(P_k)} \right) \log(1-\beta) + \text{const.}$$

Differentiating and solving gives

$$\beta = \frac{\sum_{i,g} C_{i,g}^{(M)}}{\sum_{i,g} C_{i,g}}. \tag{20}$$

**Inferring the Ambient Profile without Non-Cellular Barcodes.** For datasets with insufficient non-cellular barcodes, we assume that the ambient expression profile can be expressed as a linear combination of the cell-type expression profiles $\{\boldsymbol{p}^k\}$. In this setting, the ambient profile is treated as an additional latent mixture over cell types, and two-step training with cell-type reassignment and repulsion between $\boldsymbol{a}$ and $\{\boldsymbol{p}^k\}$ is disabled for stability.
We parameterize the ambient profile as

$$\boldsymbol{a} = \sum_{k=1}^{K} u^k \boldsymbol{p}^k, \tag{21}$$

where the mixture weights satisfy $u^k \geq 0$ and $\sum_{k=1}^{K} u^k = 1$. The mixture weights are initialized according to the estimated relative abundance of each cell type,

$$u^k = \frac{\sum_n \gamma_n^k}{\sum_{k'=1}^{K} \sum_n \gamma_n^{k'}}. \tag{22}$$

**Updating the ambient mixture weights.** The mixture weights $\{u^k\}$ are refined using a three-iteration nested EM procedure. Let $A_g$ denote the expected number of counts assigned to the ambient component feature $g$, aggregated across all barcodes.

**E-step.** Conditioned on the current estimates of $\{u^k\}$ and $\{\boldsymbol{p}^k\}$, we compute the posterior responsibility of cell type $k$ for generating ambient counts at feature $g$,

$$r_{k,g} = \mathbb{P}(k \mid g, \boldsymbol{a}) = \frac{u^k p_g^k}{\sum_{k'=1}^{K} u^{k'} p_g^{k'}} = \frac{u^k p_g^k}{a_g}. \tag{23}$$

**M-step.** The expected complete-data log-likelihood for the ambient component is

$$Q(u) = \sum_g A_g \log\left(\sum_{k=1}^{K} u^k p_g^k\right). \tag{24}$$

Maximizing $Q(u)$ with respect to $\{u^k\}$ under the simplex constraint yields the standard mixture-proportion update

$$u^k \propto \sum_g r_{k,g} A_g, \tag{25}$$

followed by normalization to enforce $\sum_k u^k = 1$.
After updating $\{u^k\}$, the ambient profile is recomputed as

$$\boldsymbol{a} \leftarrow \sum_{k=1}^{K} u^k \boldsymbol{p}^k, \tag{26}$$

.

**Stopping Conditions.** After termination of the burn-in phase, convergence is assessed using parameter-based criteria that directly measure stability of the inferred profiles and contamination fractions. Specifically, we monitor changes in the cell-type expression profiles $p_k$ and in the effective contamination fraction

$$f_i = (1-\beta)\alpha_i + \beta,$$

which captures the total non-biological contribution for each barcode.
At iteration $t$, profile convergence is quantified as

$$\Delta_p^{(t)} = \max_k \sum_{g=1}^{G} \left|p_{k,g}^{(t)} - p_{k,g}^{(t-1)}\right|,$$

corresponding to the maximum $L_1$ change across clusters. In parallel, convergence of contamination parameters is assessed using the 90$^{\text{th}}$ percentile of absolute changes in $f_i$,

$$\Delta_f^{(t)} = \text{Quantile}_{0.9}\left(\left|f_i^{(t)} - f_i^{(t-1)}\right|\right),$$

which provides a robust summary insensitive to a small number of outlier barcodes.
The algorithm is declared converged once both criteria satisfy

$$\Delta_p^{(t)} < 10^{-4} \quad \text{and} \quad \Delta_f^{(t)} < 10^{-4}.$$

These thresholds were chosen to ensure numerical stability and reproducibility across datasets, while allowing sufficient refinement of parameters after likelihood stabilization.

**Handling Cells with Extreme Ambient Fractions.** During optimization, a subset of barcodes may be driven toward extreme ambient fractions ($\alpha_i \to 1$). This behavior typically arises when a barcode is poorly explained by its assigned cell-type profile under the likelihood model. To prevent such barcodes from distorting cell-type profiles or being incorrectly explained as nearly pure ambient, CellSweep incorporates a confidence-gated reassignment mechanism.
CellSweep uses a two-stage optimization procedure:

**Stage 1: Stabilization with $\alpha$-capping.** During the initial burn-in phase (prior to convergence of the log-likelihood), per-cell ambient fractions are capped at a maximum value of 0.9. Barcodes that attempt to exceed this cap are temporarily excluded from contributing to updates of the cell-type profiles $p_k$. This ensures that poorly fit barcodes do not bias profile estimation during early optimization.

**Confidence-gated reassignment.** Only barcodes that approach the $\alpha$ cap are permitted to update their cell-type assignment during the burn-in phase. Reassignment is performed using a hard maximum-likelihood criterion over existing cell-type profiles, without allowing mixtures of cell types.
More precisely, for each barcode $i$ with observed counts $\{c_{ig}\}_{g=1}^{G}$, CellSweep re-assigns the cell-type index by maximizing the per-barcode log-likelihood under the current parameters:

$$\hat{k}_i = \arg\max_{k\in\{1,\ldots,K\}} \sum_{g=1}^{G} c_{i,g} \log\Big((1-\beta)\big((1-\alpha_i)p_{k,g} + \alpha_i a_g\big) + \beta m_g\Big). \tag{27}$$

For many barcodes, reassignment to a better-fitting profile leads to an immediate reduction in $\alpha_i$ in subsequent iterations.

**Stage 2: Final optimization.** After convergence of the log-likelihood, the $\alpha$ cap is removed and cell-type assignments are fixed. Final optimization proceeds until convergence of the model parameters.
This procedure confines cell-type reassignment to a small, well-defined subset of barcodes that would otherwise be explained as nearly pure ambient. In practice, it substantially reduces the prevalence of extreme $\alpha_i$ values while preserving overall clustering structure and stabilizing estimation of cell-type profiles.

**Output.** After convergence, the cleaned data matrix, $\widehat{\mathbf{C}}_{i,g}$ is returned.

$$\widehat{\mathbf{C}}_{i,g} = \max\{0, \mathbf{C}_{i,g} - \mathbf{C}_{i,g}^{(A)} - \mathbf{C}_{i,g}^{(M)}\}. \quad \textbf{(28)}$$

**Numerical stability.** To ensure numerical stability during inference, we apply standard safeguards to prevent undefined operations arising from zero-valued denominators or logarithms. Specifically, any denominator appearing in parameter updates is replaced by $\max(\text{denominator}, \varepsilon)$ with $\varepsilon = 10^{-12}$. Similarly, logarithms are evaluated as $\log(\max(x, \varepsilon_{\text{log}}))$, where $\varepsilon_{\text{log}} = 10^{-300}$. These thresholds are chosen to be sufficiently small so as not to affect inference in well-supported regions of the parameter space, while ensuring stable behavior in low-count regimes.

### Simulation model

To evaluate CellSweep under controlled conditions with known ground truth, we simulated scRNA-seq count matrices using a generative model that captures the dominant biological and technical features of barcode-based single-cell experiments. The simulator explicitly models discrete cell types with marker genes, shared housekeeping expression, library-size variation, barcode-specific ambient contamination, non-cellular barcodes, and global bulk contamination. The observed counts for barcode $i$ and gene $g$ are generated as

$$C_{i,g} = R_{i,g} + N_{i,g},$$

where $R_{i,g}$ denotes true biological signal and $N_{i,g}$ denotes technical noise arising from ambient and bulk contamination.

**Cell-type expression programs.** We simulate $K$ cell types over $G$ genes with cell-type proportions

$$\pi \sim \text{Dirichlet}(\mathbf{1}_K).$$

Each cell type $k$ is assigned a set of marker genes $M_k$, which are disjoint across types. Marker expression strengths are drawn from a log-normal distribution,

$$s_{k,g} \sim \text{LogNormal}(0, \sigma^2_{\text{marker}}), \quad g \in M_k,$$

and scaled to induce strong but heterogeneous marker effects. To reflect broadly expressed genes, a subset of non-marker genes is designated as housekeeping genes and assigned shared expression strengths across all cell types,

$$h_g \sim \text{LogNormal}(\mu_{\text{hk}}, \sigma^2_{\text{hk}}).$$

Each cell-type expression profile is then normalized to obtain a probability vector $p_k \in \Delta^{G-1}$. This structure produces sparse, marker-driven cell identities while retaining realistic shared expression across types.

**Cell assignments, empty droplets, and library sizes.** We simulate $N$ barcodes. Each barcode is assigned a cell type $z_i \sim \text{Categorical}(\pi)$ and independently designated as empty with probability $p_{\text{empty}}$. Non-cellular barcodes contain no biological signal ($R_{i,g} = 0$). For cellular barcodes, biological library sizes vary according to

$$L_i = L_0 \exp(\epsilon_i), \quad \epsilon_i \sim \mathcal{N}(\mu_L, \sigma_L^2),$$

reflecting variability in capture efficiency and sequencing depth.

**Biological count generation.** For cell-containing barcodes, biological counts are generated using a Gamma–Poisson construction to introduce overdispersion:

$$\lambda_{i,g} \sim \text{Gamma}\left(r, \frac{L_i p_{z_i,g}}{r}\right), \qquad R_{i,g} \sim \text{Poisson}(\lambda_{i,g}),$$

where $r$ controls dispersion. This yields negative-binomial-like variability in gene expression while preserving the expected cell-type structure.

**Ambient contamination.** Ambient RNA is modeled as a shared gene distribution reflecting the population-level transcriptome. We define the ambient profile as

$$a = (1 - \lambda) \sum_{t=1}^{K} \pi_t p_t + \lambda u,$$

where $u$ is a diffuse background distribution and $\lambda$ controls low-level leakage into all genes. This construction captures the empirical observation that ambient RNA resembles a mixture of cellular expression programs rather than an arbitrary background.
For each barcode $i$, the total ambient load is drawn from a log-normal distribution,

$$\Lambda_i \sim \text{LogNormal}(\mu_\alpha, \sigma_\alpha^2),$$

with larger variability for empty droplets than for cell-containing droplets. Conditional on $\Lambda_i$, ambient counts are sampled as

$$N_{i,g}^{(\text{amb})} \sim \text{Poisson}(\Lambda_i a_g).$$

This produces substantial heterogeneity in ambient fractions across barcodes, including cells with very low contamination and others dominated by background.

**Global bulk contamination.** To model contamination introduced after barcode assignment (e.g., barcode swapping or index hopping), we introduce a global bulk noise process. A fraction $\beta$ of total molecules is redistributed across barcodes while preserving gene identity. This produces a low-rank, approximately uniform background signal shared across cells, which cannot be explained by droplet-specific ambient capture alone.

**Ground-truth annotation.** The simulator records the observed counts $C_{i,g}$, the true biological signal $R_{i,g}$, and the injected noise $N_{i,g}$ separately. The realized per-barcode ambient fraction

$$\alpha_i = \frac{\sum_g N_{i,g}}{\sum_g C_{i,g}}$$

is reported for non-empty barcodes, with $\alpha_i = 1$ for empty droplets. These ground-truth quantities enable direct assessment of background-removal accuracy, signal preservation, and robustness across contamination regimes.
